# Multi-site Cleavage of Amyloid-β by a Minimal 5-mer Catalytic Peptide: Mimicking Serine Protease Activity via Hydrophobic Anchoring

**DOI:** 10.64898/2026.06.03.729455

**Authors:** Fumiaki Ito, Motomi Konishi, Rina Nakamura, Toshifumi Akizawa

**Affiliations:** The Institute of Prophylactic Pharmacology, 1-58, Rinku-oraikita, Izumisano 598-8531, Osaka, Japan; Department of Integrative Pharmacy, Faculty of Pharmaceutical Sciences, Setsunan University, 45-1 Nagaotoge-cho, Hirakata 573-0101, Osaka, Japan; O-Force Co., Ltd., 3454 Irino Kuroshio-cho, Hata-gun 789-1931, Kochi, Japan; Department of Pharmacology, Kochi Medical School, Kochi University, Kohasu, Oko-cho, Nankoku 783-8505, Kochi, Japan

**Keywords:** Alzheimer’s disease, amyloid-*β*, neurodegenerative disease, catalytides, catalytic peptides, peptide drug, serine proteases, amyloid-*β* cleavage

## Abstract

The development of small synthetic catalytic peptides, or ”catalytides,” offers a promising therapeutic strategy for the targeted degradation of amyloid-beta (A*β*). Among these, the pentapeptide SKGQA mimics the proteolytic activity of serine proteases despite its minimal size. However, the molecular mechanism enabling such a short, highly flexible peptide to execute cleavage across multiple sites remains unclear. In this study, we combined HADDOCK docking and extended molecular dynamics (MD) simulations to investigate the dynamic interaction between SKGQA and the central hydrophobic cluster (CHC, residues 17–21) of A*β*. Our MD simulations reveal a cooperative mechanism underlying this activity. The hydrophobic C-terminal Ala5 residue anchors SKGQA within the A*β* CHC region, stabilizing the bound complex while potentially interfering with A*β* self-assembly through steric hindrance. Simultaneously, while the C-terminus remains anchored, the hydrophilic N-terminal Ser1 residue is positioned in spatial proximity to multiple adjacent backbone carbonyl carbons—including Glu22 and Asp23. Within this anchored conformational ensemble, thermal fluctuations allow Ser1 to stochastically attack these neighboring carbonyl targets. This combination of site-specific hydrophobic anchoring and probabilistic catalytic targeting provides a clear structural rationale for the peptide’s multi-site cleavage capability.

## 1 Introduction

The accumulation of amyloid-*β* (A*β*) 42, particularly in its oligomeric and fibrillar forms, is a central pathological event in Alzheimer’s disease (AD)[1, 2]. While recent advancements in monoclonal antibodies such as lecanemab have demonstrated clinical efficacy in clearing A*β* plaques, challenges regarding high costs, delivery efficiency, and safety concerns (e.g., amyloid-related imaging abnormalities [ARIA] involving brain edema) remain[3, 4]. As an alternative, we have focused on “catalytides”—short synthetic proteolytic peptides that endogenously degrade A*β*[5, 6]. Among these, SKGQA, a 5-mer peptide derived from the ANA/BTG3 protein, represents one of the smallest functional units capable of protease-like activity[7].

A defining characteristic of classical serine proteases is the “catalytic triad,” typically composed of Ser, His, and Asp[8], which activates the Ser hydroxyl group for nucleophilic attack. Previously, we proposed that even minimal catalytides like SKGQA can satisfy these functional requirements by forming an intramolecular pseudo-catalytic triad, wherein the N-terminal amine (–NH_2_) acts as a base, the C-terminal carboxylate (– COOH) stabilizes the conformation, and the Ser1 hydroxyl group (–OH) serves as the nucleophile[7].

Building upon this concept, we hypothesized that transient intramolecular N–C terminal proximity stabilizes SKGQA in a compact “reactive conformation,” positioning the Ser1 hydroxyl group in a spatially favorable orientation for nucleophilic attack on the substrate backbone.

Crucially, experimental evidence indicates that SKGQA does not target a single conserved sequence but instead cleaves both soluble and solid-state A*β* at multiple, diverse sites[7]. This multi-site cleavage suggests a unique structural flexibility, allowing the 5-mer to adapt its conformation to various regions of the A*β* molecule. However, how such a short peptide transiently stabilizes this pseudo-catalytic geometry while maintaining the dynamic adaptability required for non-specific targeting across substrate environments remains to be fully elucidated.

In this study, we investigate the molecular mechanism underlying the multi-site proteolytic action of SKGQA. To provide a structural rationale for its cleavage capability, we characterize the dynamic ensemble of SKGQA–A*β*_17–42_ complexes using HADDOCK docking and molecular dynamics (MD) simulations. Our findings reveal a cooperative mechanism in which site-specific hydrophobic anchoring within the central hydrophobic core (CHC) stabilizes the bound complex, while the intrinsic flexibility of Ser1 enables stochastic sampling of adjacent backbone carbonyl targets. This dynamic balance between localized anchoring and probabilistic targeting provides new insights into the design of adaptable, low-molecular-weight catalytides.

## 2 Methods

### 2.1 Initial Structure Prediction using PEP-FOLD3

Initial three-dimensional structures of the peptides were predicted from their primary amino acid sequences using the PEP-FOLD3 web server (https://bioserv.rpbs.univ-paris-diderot.fr/services/PEP-FOLD3/). PEP-FOLD3 is a *de novo* computational approach designed for the structural prediction of peptides[9, 10, 11]. From the simulation results, seven candidate models (Models 1–7) were obtained in PDB format for further analysis.

### 2.2 Constraint-based Modeling with MODELLER

To design the triad-forming peptides, structural refinement was performed using MODELLER 10.6, a software package for protein structure homology modeling and prediction [12]. Model 1, obtained from PEP-FOLD3, was utilized as the structural template. Specific spatial constraints were applied to the N-terminal Serine (Ser) and C-terminal Alanine (Ala) residues. Using the MODELLER script, we generated a structural ensemble where the distance between the N-terminal and C-terminal residues was constrained to 3.5 Å to facilitate triad formation. This moderate constraint was chosen to avoid artificial steric strain prior to docking, enabling unrestrained dynamic sampling during subsequent MD simulations. A total of 10 refined models (Target 1–10) were generated, corresponding to the output files target.B99990001.pdb through target.B99990010.pdb. Among the generated ensemble, Target 1 (target.B99990001.pdb; N–C distance of 3.7 Å) was selected as the representative initial conformation for subsequent molecular docking.

### 2.3 Structural Analysis and Visualization

Structural visualization and geometric measurements were conducted using the open-source version of PyMOL (version 2.6, Schrödinger, LLC)[13]. The software environment was configured on a Windows platform by installing Python 3 and the Microsoft Visual C++ Redistributable for Visual Studio.

The distance between the N-terminal Ser and C-terminal Ala of the SKGQA sequence in each of the ten target models (Target 1–10) was measured and verified using PyMOL to ensure compliance with the design constraints.

### 2.4 Molecular Docking using HADDOCK

Molecular docking was performed to investigate the binding interactions between a single monomeric chain derived from the A*β*_17–42_ structure (PDB ID: 2BEG [14]) and the SKGQA peptide using the HADDOCK web server, version 2.4 [15, 16]. The docking process was guided by defining active residues: residues 17–21 (LVFFA) were designated as the active residues for the A*β* target region, while all residues of the SKGQA peptide were defined as active for the catalytide. Following the standard HADDOCK protocol, docking parameters allowed for flexible interface refinement. Generated poses were clustered based on structural similarity (root-mean-square deviation, RMSD) and energetic characteristics. The simulation yielded multiple docking clusters; in this study, we focused on Clusters 1 and 7, which exhibited the most favorable HADDOCK scores. Specifically, the top four structures from these clusters (e.g., Cluster 1-1 to 1-4) were subjected to detailed structural and energetic analysis, as they represented the most energetically favorable and stable binding modes within the ensemble.

### 2.5 Molecular Dynamics Simulations and Trajectory Analysis

Molecular dynamics (MD) simulations were performed using OpenMM (version 8.1) [17] with the AMBER14SB force field [18]. The molecular complex formed by a single monomeric chain of A*β*_17–42_ and the SKGQA peptide was explicitly solvated in a cubic TIP3P water box [19] extending 1.0 nm from the solute boundaries, and neutralized with 0.15 M NaCl. Long-range electrostatic interactions were treated using the smooth particle mesh Ewald (PME) method [20] with a non-bonded cutoff of 1.0 nm. All peptide and protein N- and C-termini were modeled in their uncapped, zwitterionic states (NH^+^ and COO*^−^*) at physiological pH 7.4, with standard protonation states assigned to all ionizable residues.

Following energy minimization (L-BFGS algorithm, maximum force *<* 10.0 kJ/mol/nm) and two-stage equilibration (NVT at 310 K for 10 ps, followed by NPT at 1.0 atm for 10 ps), production runs were executed.

To systematically evaluate the dynamic interactions, both short-term (2 ns) and long-term (50 ns, 3 runs) production simulations were conducted. For the 50-ns production runs, three independent trajectories were generated using randomized initial atomic velocity assignments to ensure sampling breadth and statistical independence. Structural snapshots were recorded every 10 ps.

Trajectory analyses were performed using MDTraj [21] and custom Python scripts. To evaluate overall structural stability, C*α* RMSD profiles were calculated relative to the initial configuration (*t* = 0 ns, frame 0) across the complete 50-ns production trajectories (*n* = 3). To further characterize local binding consistency, ensemble-averaged alignment RMSDs were determined for bound states, defined by filtering frames where the minimum inter-molecular distance between SKGQA and A*β*_17–25_ was *≤* 5.0 Å. Furthermore, to evaluate catalytic proximity and internal peptide dynamics, inter-atomic distance trajectories were monitored across the simulations, focusing specifically on the backbone N–C distance within SKGQA (Ser1 to Ala5) and the nucleophilic distances between Ser1(O*γ*) and the backbone carbonyl carbons of the A*β*_17–25_ region.

### 2.6 Contact Mapping and Representative Three-Dimensional Binding Geometry

To capture the comprehensive interaction profile within the reaction region of the substrate, inter-residue contact dynamics between the A*β*_17–25_ region (of the A*β*_17–42_ monomer) and SKGQA were analyzed. Two-dimensional contact maps were generated based on the average minimum heavy-atom distances under defined analytical criteria: the overall 50-ns ensemble, the bound state (inter-peptide distance *≤* 5.0 Å), and the reactive state (Ser1-O*γ* to A*β* backbone C=O distance *≤* 4.0 Å). Structural clustering was applied to identify representative centroid structures from the trajectories for detailed characterization of residue-residue interactions. Furthermore, representative three-dimensional binding geometries and atomic-level catalytic interactions in the reactive state were extracted and visualized to clarify the spatial orientation and nucleophilic attack pathways of the SKGQA peptide relative to the substrate core.

## 3 Results

### 3.1 Conformational Flexibility of SKGQA *in silico*

To elucidate the structural basis for the proteolytic activity of the 5-mer peptide SKGQA, we initially analyzed its conformational space using PEP-FOLD structure prediction. We previously hypothesized that the catalytic activity of SKGQA is mediated by a triad-like arrangement involving the Ser hydroxyl group, the N-terminal amino group (-NH_2_), and the C-terminal carboxylate (-COOH)[7].

The *de novo* modeling results, summarized in Table 1, revealed that SKGQA predominantly exists in a range of flexible, non-catalytic conformations. The calculated sOPEP (Simplified Protein Energy Potential) values for the top-ranked models were nearly indistinguishable (ranging from approximately -1.37 to -1.33), suggesting the absence of a uniquely stable, energy-minimized fold.

**Table 1:** Predicted energy and inter-atomic distances of SKGQA models by PEP-FOLD.

| Model | sOPEP Energy (a.u.) | Ser($O\gamma$)-Ser(N) (Å) | Ser(N)-Ala(O1) (Å) | Ser(N)-Ala(O2) (Å) |
| --- | --- | --- | --- | --- |
| 1 | -1.370 | 2.6 | 7.0 | 5.4 |
| 2 | -1.370 | 2.6 | 7.1 | 5.4 |
| 3 | -1.370 | 2.6 | 7.1 | 5.4 |
| 4 | -1.370 | 2.6 | 7.0 | 5.4 |
| 5 | -1.355 | 2.6 | 7.3 | 5.6 |
| 6 | -1.370 | 2.6 | 7.0 | 5.4 |
| 7 | -1.328 | 2.7 | 8.4 | 6.4 |

*Note:* a.u.: arbitrary units. The sOPEP score in PEP-FOLD3 represents a coarse-grained energy function in dimensionless units. Atom designations O1 and O2 refer to the two oxygen atoms of the C-terminal carboxyl group.

Crucially, in the representative models generated by PEP-FOLD, the spatial distances required for triad formation were not satisfied. While the distance between the Ser hydroxyl group and the N-terminal amino group (O*γ*-N) was relatively close (approximately 2.6-2.7 Å), the distances between the N-terminal nitrogen (N) and the C-terminal carboxyl oxygens of Alanine (O1 and O2) were consistently greater than 5 Å (Table 1). These *in silico* models represent the most probable conformations in an isolated, substrate-free state, indicating that SKGQA does not spontaneously form a stable catalytic triad under these conditions.

### 3.2 Construction of Triad-like Active Models using MODELLER

While the majority of isolated conformations are inactive, we hypothesized that the peptide could transiently adopt or be induced into a triad-like arrangement favorable for catalysis. To explore the structural feasibility of such a state, we investigated the candidate structures generated by PEP-FOLD (Models 1–7). As Models 1–7 shared comparable energetic and geometric features, Model 1 was selected as a representative starting template for further structural refinement.

Comparative modeling was performed using MODELLER to generate 10 target models by imposing specific distance restraints between the N-terminal nitrogen and the C-terminal carboxyl oxygen. Across the 10 generated target models, the N–C distance was consistently maintained within a narrow range of 3.5 to 3.7 Å. From this ensemble, Target 1 (target.B99990001.pdb; N–C distance of 3.7 Å) was selected as the representative initial conformation for subsequent molecular docking. As shown in Figure 1, the induced active model exhibited a significantly more compact, ”curled” conformation compared to the relaxed structure of the initial PEP-FOLD Model 1. This structural transition demonstrates that the 5-mer peptide can satisfy the geometric requirements for a serine protease-like catalytic triad through relatively minor conformational adjustments.

**Figure 1:**
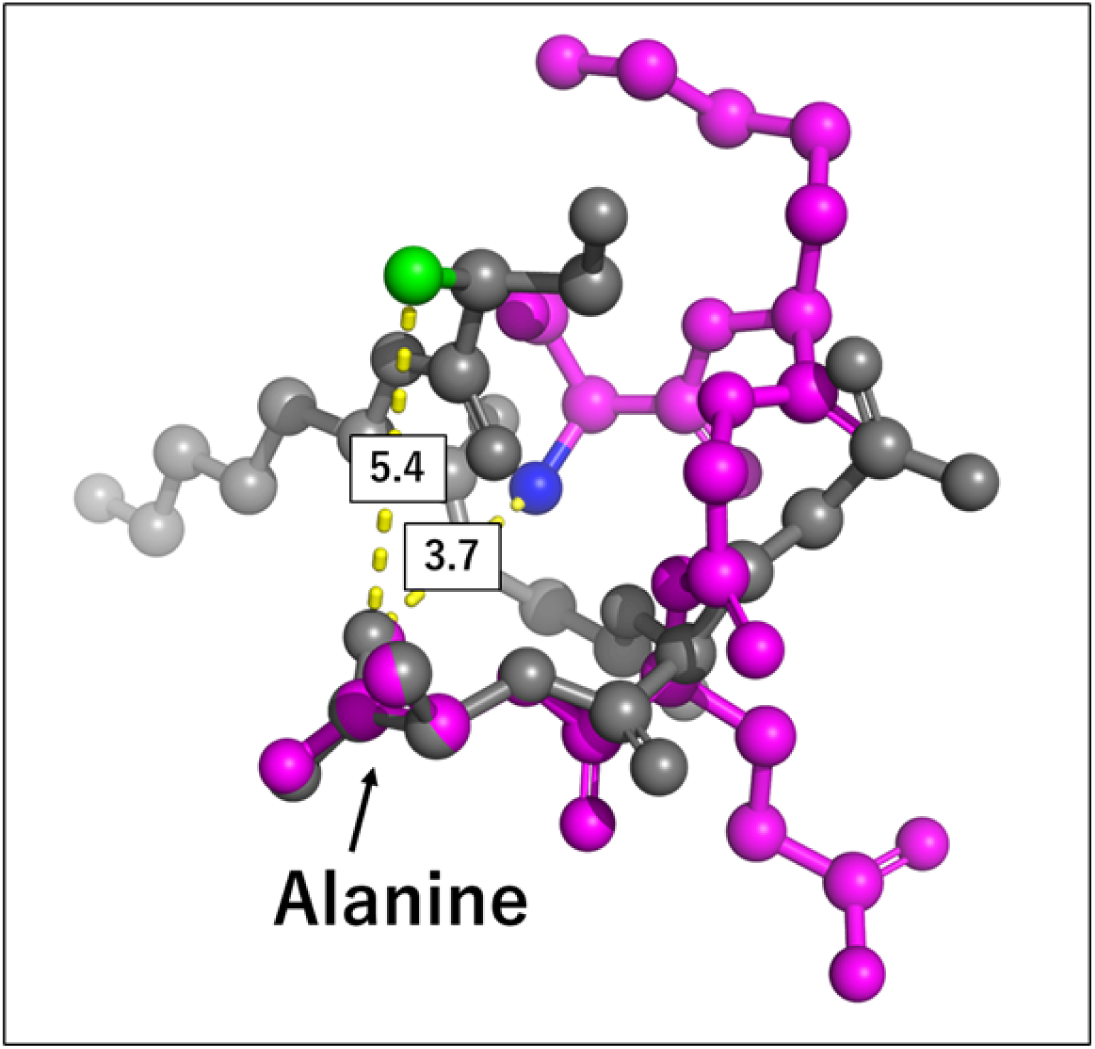
Structural comparison of SKGQA between the predicted and induced active states. The *de novo* structure predicted by PEP-FOLD (gray) shows a relaxed conformation with a large distance (5.4 Å) between the N-terminal amino group and the C-terminal carboxyl group. In contrast, the active model induced by MODELLER (magenta) demonstrates a constrained, curled conformation where these functional groups are brought into proximity (3.7 Å). The structures were aligned based on the fifth alanine residue (indicated by arrow). Distances are indicated by yellow dashed lines (unit: Å).

### 3.3 HADDOCK Docking Analysis and Binding Conformational Diversity

To explore the initial interaction and binding modes between the active SKGQA model (Target 1) and A*β*, molecular docking simulations were performed using the HADDOCK server. The A-chain of the monomeric A*β*_17–42_ structure derived from the fibril model (PDB ID: 2BEG [14]) was selected as the receptor. The active target binding site was defined as the central hydrophobic cluster (CHC, residues 17–21: LVFFA) (Figure 2).

**Figure 2:**
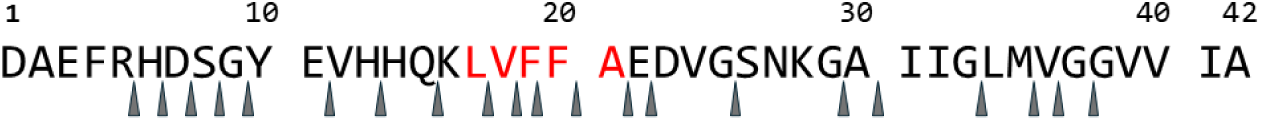
The amino acid sequence of A*β*_1–42_ and its major target cleavage sites (indicated by arrows). The central hydrophobic cluster (CHC) sequence 17–21 (LVFFA), highlighted in red, was designated as the active target region for docking simulations.

The HADDOCK clustering analysis categorized 153 generated complex structures into 13 distinct clusters. To perform a comprehensive and unbiased structural screening, we utilized PyMOL to systematically examine the binding modes, internal geometries, and energetic criteria across the clusters. Based on HADDOCK scores and favorable *Z*-scores, Cluster 1 and Cluster 7 were identified as the primary functional clusters. All top-ranking models within these clusters—namely Clusters 1-1, 1-2, 1-3, and 1-4 from Cluster 1, and Clusters 7-1, 7-2, 7-3, and 7-4 from Cluster 7—were comprehensively analyzed (Figure 3).

**Figure 3:**
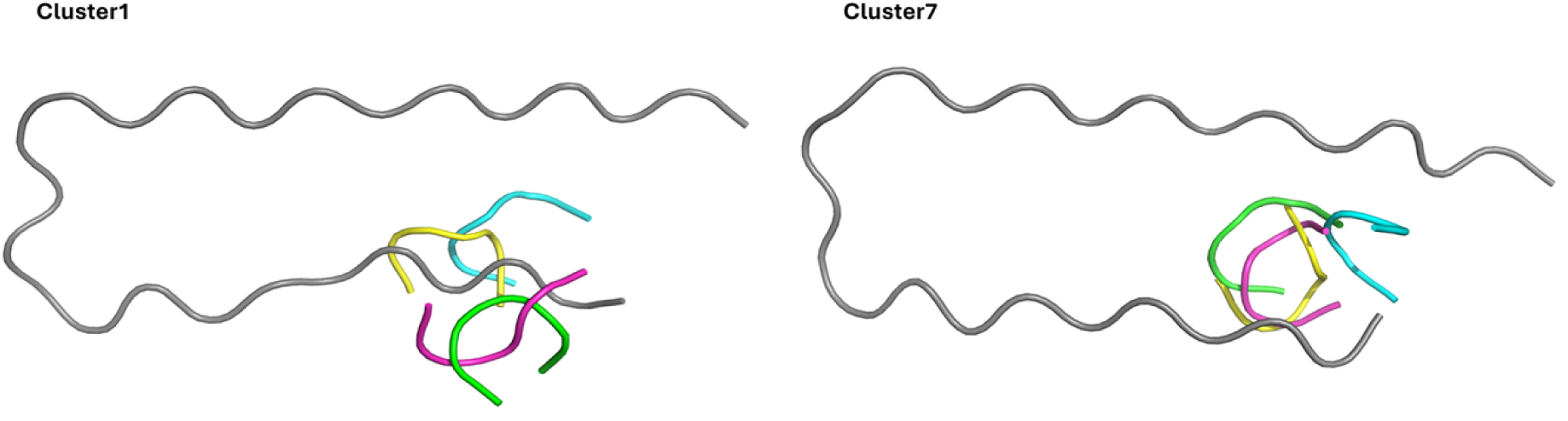
Superimposed binding modes of SKGQA on A*β*_17–42_ derived from HADDOCK docking clusters. Structural overlay of the top four models from Cluster 1 (left) and Cluster 7 (right) bound to the A*β*_17–42_ receptor backbone (shown in gray tube representation). In both panels, individual SKGQA models are color-coded as follows: green for Cluster 1 (1-1 or 7-1), cyan/blue for cluster 2 (1-2 or 7-2), magenta for Cluster 3 (1-3 or 7-3), and yellow for Cluster 4 (1-4 or 7-4).

As shown in Figure 3, SKGQA binds predominantly near the N-terminal segment of A*β*_17–42_—specifically in the vicinity of the central hydrophobic cluster (CHC, residues 17–21)—adopting diverse binding orientations rather than a single rigid conformation. Detailed structural examination of the bound SKGQA peptides revealed significant conformational flexibility upon interaction with A*β*_17–42_ (Figures S1 and S2). While the uncomplexed SKGQA structure used as the initial docking input exhibited an intramolecular Ser1(N) to Ala5(O) distance of 3.7 Å, the corresponding distances within the bound models varied substantially, ranging from 4.3 Å to 8.9 Å in Cluster 1 and 4.6 Å to 8.5 Å in Cluster 7 (Table 2).

**Table 2:**
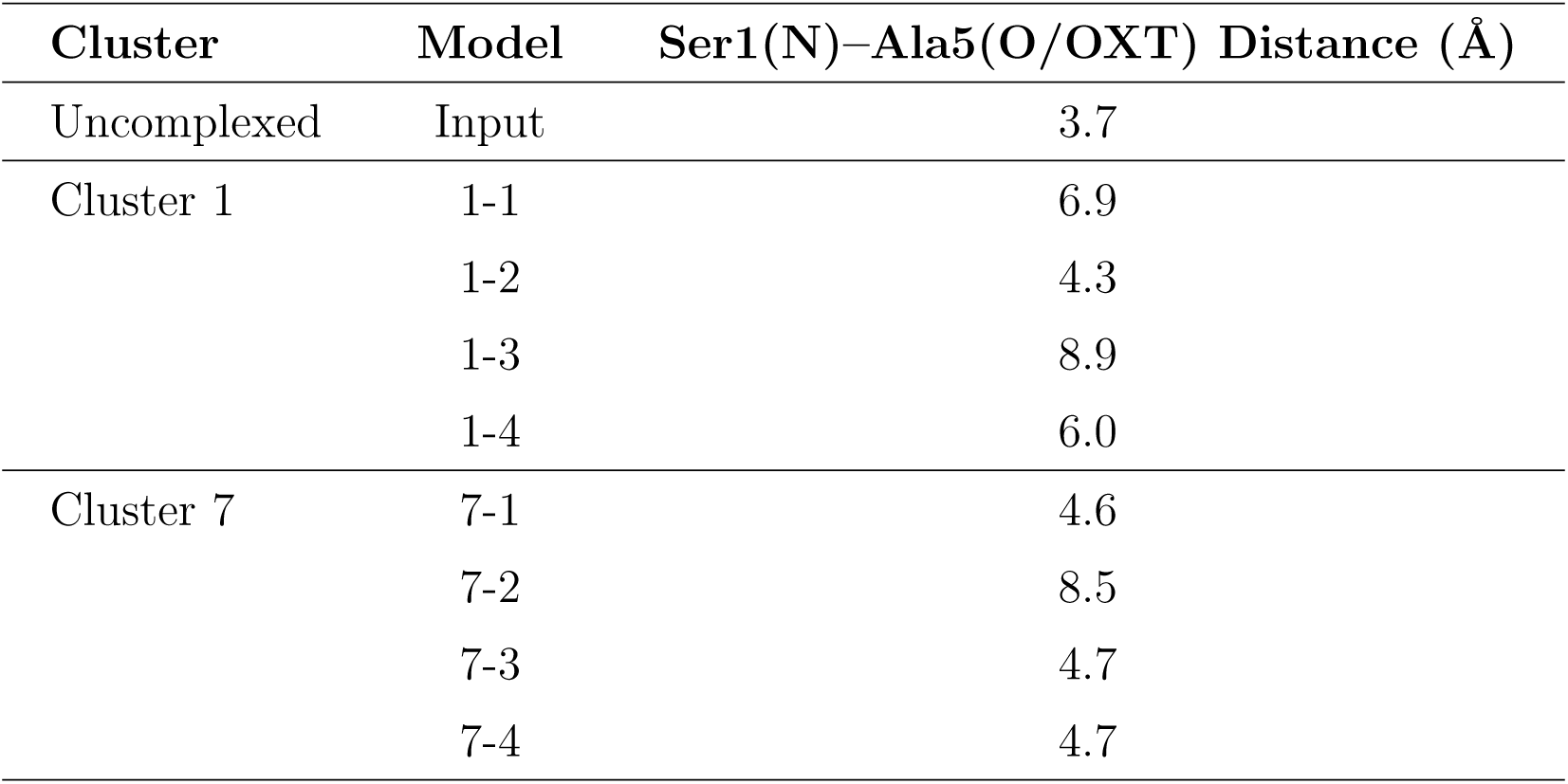
Intramolecular Ser1(N)–Ala5(O/OXT) distances in the uncomplexed and HADDOCK-docked SKGQA models.

| Cluster | Model | Ser1(N)–Ala5(O/OXT) Distance (Å) |
| --- | --- | --- |
| Uncomplexed | Input | 3.7 |
| Cluster 1 | 1-1 | 6.9 |
|  | 1-2 | 4.3 |
|  | 1-3 | 8.9 |
|  | 1-4 | 6.0 |
| Cluster 7 | 7-1 | 4.6 |
|  | 7-2 | 8.5 |
|  | 7-3 | 4.7 |
|  | 7-4 | 4.7 |

These structural variations demonstrate that SKGQA undergoes dynamic conformational adaptation upon engaging A*β*_17–42_, resulting in a heterogeneous ensemble of initial complexes that serve as starting points for subsequent molecular dynamics simulations.

### 3.4 Short-Term Molecular Dynamics Simulations and Selection of Representative Candidates

To evaluate the dynamic stability of the initial docking poses under thermodynamic fluctuations, short-term (2 ns) explicit-solvent molecular dynamics (MD) simulations were conducted for all models within Cluster 1 (1-1 to 1-4) and Cluster 7 (7-1 to 7-4). As shown in Figure 4, using the final structure at *t* = 2.0 ns as the reference frame, the backbone C*α* RMSD values for all evaluated clusters gradually decreased from initial peak values (up to 11–12 Å) toward the reference bound state. In Cluster 1, Cluster 1-1 and Cluster 1-4 exhibited rapid structural relaxation toward stable binding poses, with RMSD values dropping below 4 Å after 1.0 ns and smoothly reaching equilibrium near *t* = 2.0 ns. Within Cluster 7, Cluster 7-1 and Cluster 7-3 demonstrated similarly steady equilibration trajectories, consistently progressing toward stable reference structures over the simulation period.

**Figure 4:**
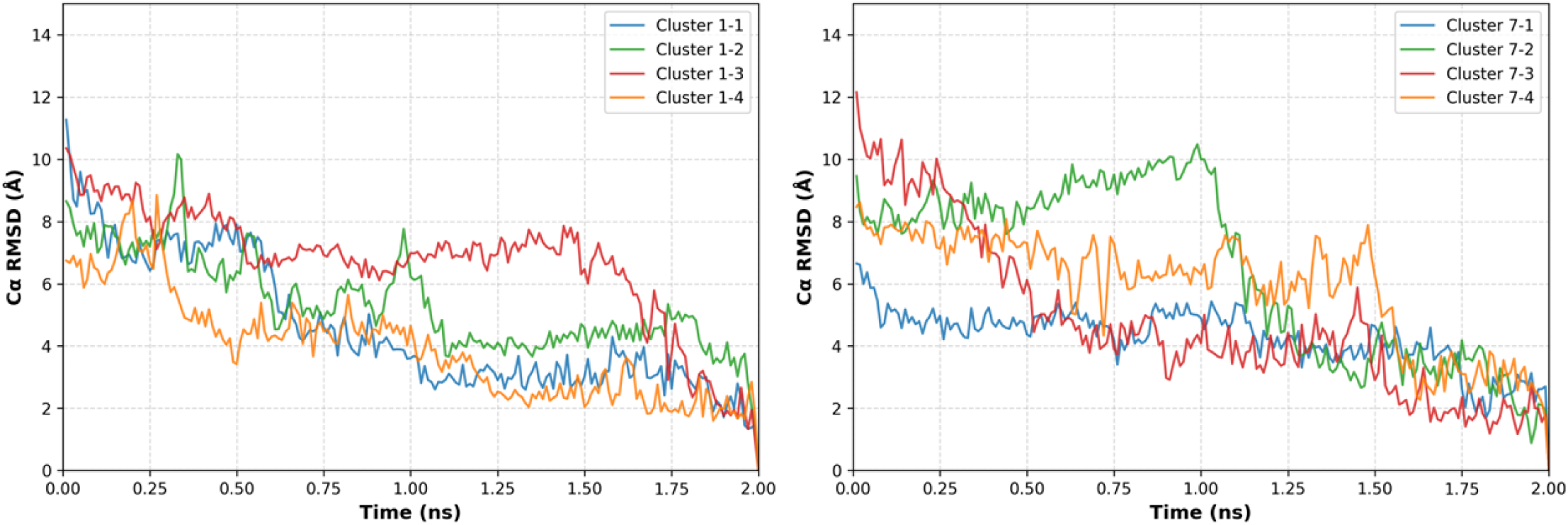
Time-dependent C*α* RMSD profiles of the SKGQA/A*β*_17–21_ complex interfaces during initial relaxation. The values calculated for the C*α* atoms of the binding interface—comprising A*β*_17–21_ (Leu17–Ala21) and the bound SKGQA peptide—are plotted over the 2.0-ns simulation period using the final structure at *t* = 2.0 ns as the reference frame to track structural relaxation and binding ensemble stability. The left panel displays the trajectories for Cluster 1-1, 1-2, 1-3, and 1-4, while the right panel displays those for Cluster 7-1, 7-2, 7-3, and 7-4. Individual structures within each cluster are color-coded as shown in the inset legends: blue (-1), green (-2), red (-3), and orange (-4).

To assess the conformational integrity relevant to catalytic activity, the terminal distance within the bound SKGQA peptide was monitored (Figure 5). Although temporal fluctuations were observed across all clusters—reflecting the intrinsic flexibility of the non-covalent complex—several clusters periodically adopted compact orientations where the distance approached the catalytic threshold (*<* 4 Å). Specifically, during the second half of the 2-ns trajectories, Clusters 1-1, 1-3, 1-4, and 7-2 displayed frequent transient approaches to this close proximity.

**Figure 5:**
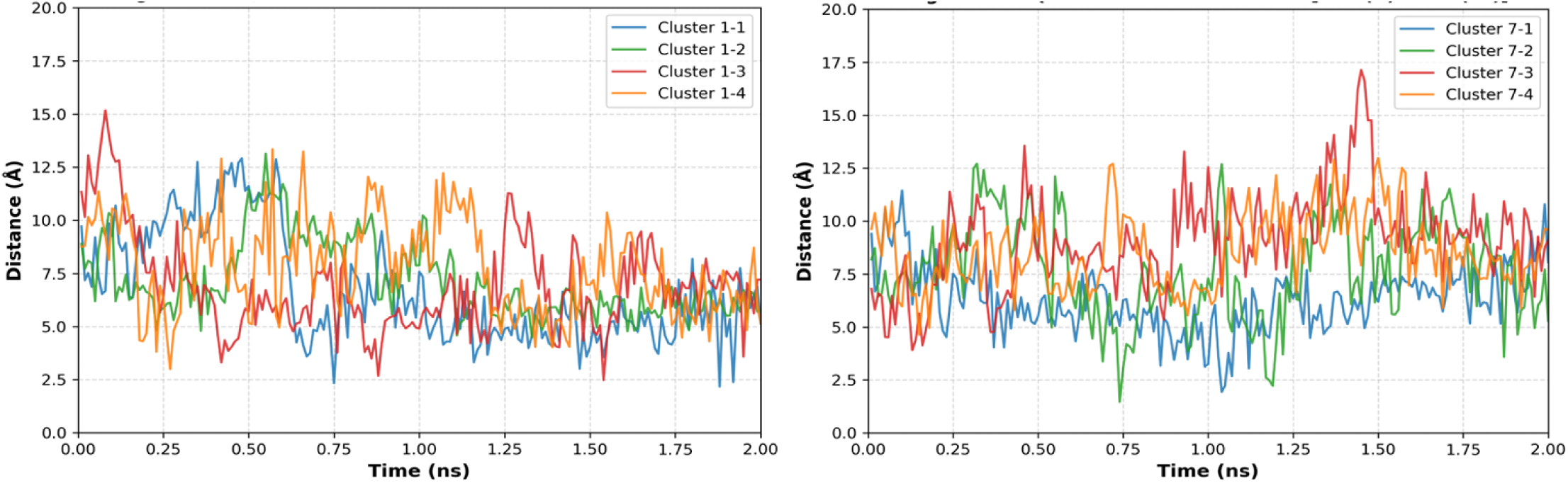
Time-dependent intramolecular Ser1(N)–Ala5(O/OXT) distance profiles of bound SKGQA. The distance between the N-terminal nitrogen atom of Ser1 [Ser1(N)] and the C-terminal carboxylate oxygen of Ala5 [Ala5(O/OXT)] within the bound SKGQA peptide is plotted over the 2.0-ns simulation period. The left panel displays the trajectories for Cluster 1-1, 1-2, 1-3, and 1-4, while the right panel displays those for Cluster 7-1, 7-2, 7-3, and 7-4. Individual structures within each cluster are color-coded as shown in the inset legends: blue (-1), green (-2), red (-3), and orange (-4).

Based on the combined criteria of lower RMSD values relative to the bound target (structural stability) and short terminal/interatomic distances (potential for reactive geometry), Clusters 1-1 and 1-4 were selected as representative candidate clusters for extended long-term (50 ns) MD simulations to further investigate their conformational dynamics and binding persistence.

### 3.5 Extended Long-Term Dynamics and Catalytic Proximity of Selected Candidates

To achieve statistical robustness and capture large-scale conformational sampling, extended 50-ns MD simulations were performed in triplicate (*n* = 3) for both Cluster 1-1 and Cluster 1-4. Figure 6 presents the Global trajectory Cα RMSD, intramolecular N–C distances within SKGQA (Ser1-N to Ala5-O*δ*1 and Ala5-O*δ*2), and the catalytic distance between the nucleophilic Ser1(O*γ*) of SKGQA and the backbone carbonyl carbons (*C* = O) of the A*β*_17_*_−_*_21_ segment.

**Figure 6:**
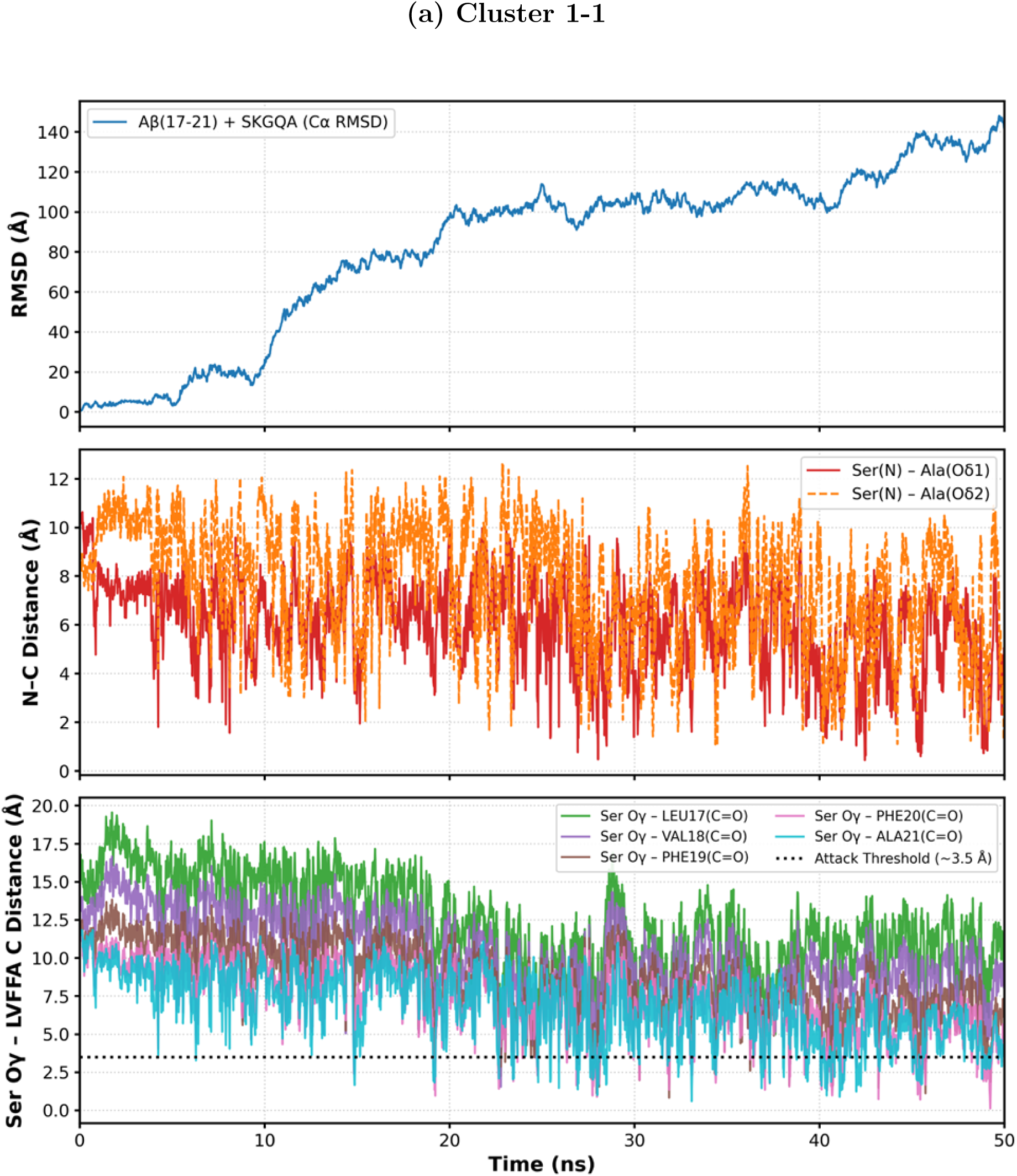
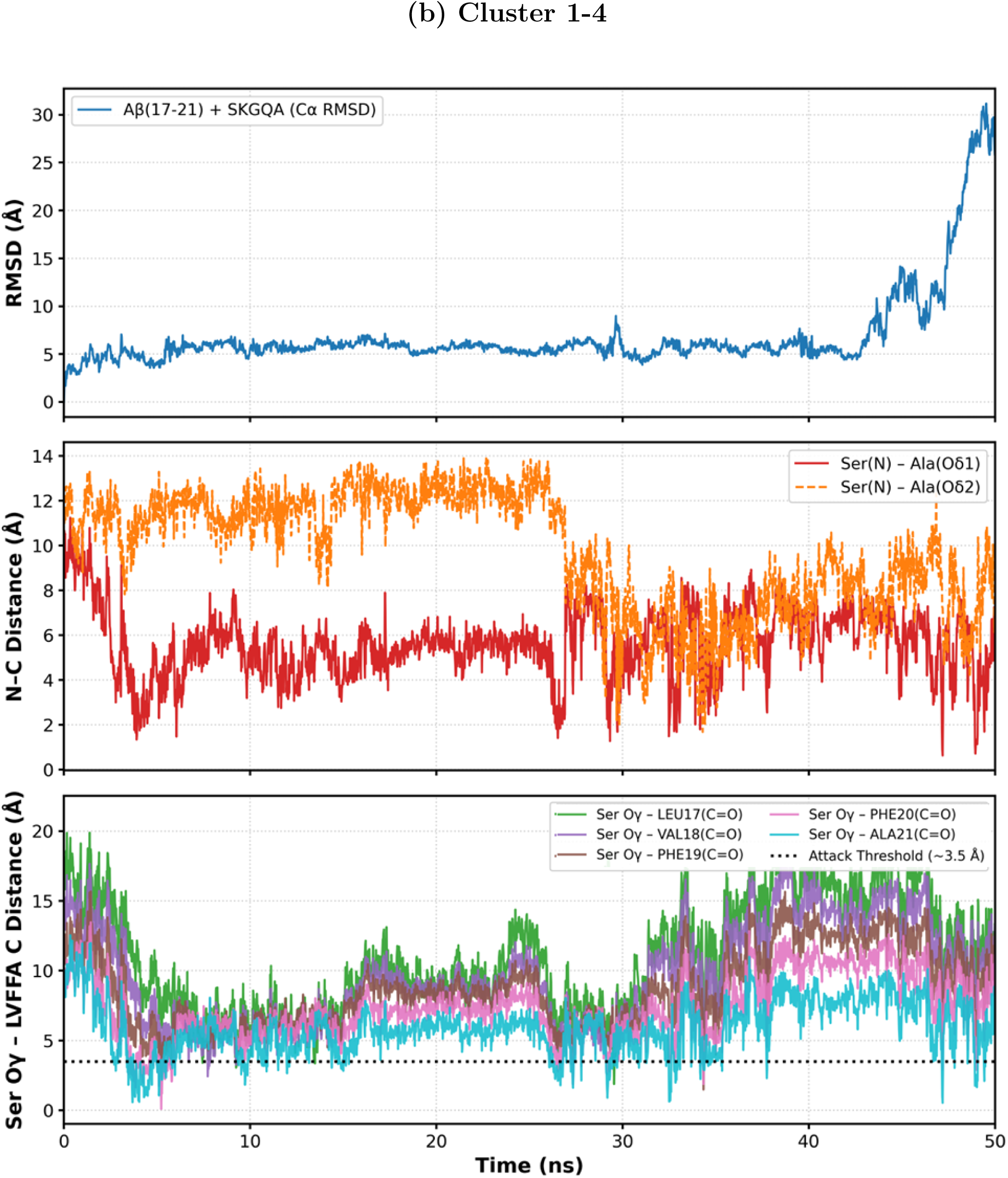
Comparative 50-ns molecular dynamics trajectory analyses of Cluster 1-1 (a) and Cluster 1-4 (b) complexes. Trajectory profiles over the 0–50 ns simulation period are shown for each complex. **Top panels:** Global trajectory C*α* RMSD (Å) of the A*β*_17–21_–SKGQA complex interface, presented as mean *±* SD (Cluster 1-1: 134.44 *±* 61.91 Å; Cluster 1-4: 11.66 *±* 7.92 Å). **Middle panels:** Intramolecular N–C distance trajectories (Å) within SKGQA, specifically monitoring the distances between Ser1(N) and the terminal carboxylate oxygens, Ala5(O*δ*1) (red solid line) and Ala5(O*δ*2) (orange dashed line). **Bottom panels:** Interatomic distances (Å) between the Ser1 nucleophilic oxygen (Ser O*γ*) and the backbone carbonyl carbons (C=O) of A*β*_17–21_ (Leu17–Ala21). The dotted horizontal black line indicates the reactive threshold for nucleophilic attack (*∼* 3.5 Å).

For Cluster 1-1 (Figure 6a), the complex C*α* RMSD remained low during the initial 2-ns phase, but exhibited a marked increase, reaching approximately 100–140 Å over the 50-ns timescale. This large RMSD elevation reflects major translational or rotational reorientations relative to the initial binding pose, indicating high conformational mobility or transient partial detachment from A*β*. Nevertheless, periodic close approaches between Ser1(O*γ*) and the backbone carbonyl carbons (*C* = O) of A*β*_17–21_ were observed throughout the trajectory, frequently crossing the spatial threshold (*∼* 3.5 Å) required for nucleophilic attack. This catalytic proximity was further facilitated by the internal dynamics of SKGQA, where the intramolecular N–C distance periodically contracted to below 2 Å, temporarily stabilizing a compact conformation conducive to reactive engagement.

Conversely, for Cluster 1-4 (Figure 6b, top panel), the complex C*α* RMSD remained low throughout the majority of the 50-ns simulation, maintaining a persistent global binding mode. The transient RMSD elevation observed toward the end of the trajectory (*>* 45 ns) reflects the inherent thermodynamic flexibility and partial transient unbinding characteristic of short, flexible catalytides. Despite these minor unbinding events, Cluster 1-4 maintained a significantly lower overall C*α* RMSD (11.66 *±* 7.92 Å) compared to Cluster 1-1 (134.44 *±* 61.91 Å). This thermodynamic stability directly correlated with high catalytic accessibility: as shown in Figure 6b (bottom panel), Ser1(O*γ*) maintained persistent, close approaches to the backbone carbonyl carbons (*C* = O) of A*β*_17–21_, frequently and continuously remaining below the reactive threshold (*∼* 3.5 Å) between 3 ns and 35 ns. Concurrently, the intramolecular N–C distances within SKGQA (Figure 6b, middle panel) exhibited active fluctuations, with both Ser1(N)–Ala5(O*δ*1) and Ser1(N)–Ala5(O*δ*2) distances periodically dropping below 2 Å. These continuous dynamics indicate that rather than rigidifying into a single conformer, Cluster 1-4 operates in a dynamic equilibrium that maintains reactive catalytic proximity while accommodating conformational flexibility despite the intrinsic mobility of short peptides.

To confirm the statistical consistency underlying these ensemble averages (*n* = 3), individual trajectories (Run 1, Run 2, and Run 3) were examined separately (Figures S3 and S4). Although each run displayed distinct temporal fluctuations in the N–C distance, all three independent simulations exhibited consistent overall trends, supporting the reproducibility of the dynamic ensemble behavior for both Cluster 1-1 and Cluster 1-4. Taken together, these MD trajectory analyses confirm that while short catalytides naturally sample a flexible conformational landscape, Cluster 1-4 effectively stabilizes the complex and maintains the catalytic Ser1 residue within productive attack distance of the A*β* backbone.

### 3.6 Inter-molecular Contact Map Analysis and Spatial Distribution in Bound States

To clarify the physical interactions governing complex formation, residue-level contact maps were constructed for the A*β*_17–25_ region over the 50-ns simulation trajectories. As shown in Supplementary Figure S5, the time-averaged minimum inter-residue distances (calculated across all frames) revealed a stark contrast between Cluster 1-1 and Cluster 1-4. In Cluster 1-1, these average distances were extremely large (*>* 200 Å), reflecting a predominantly dissociated state where the SKGQA peptide diffuses freely away from A*β*_17–25_. In contrast, Cluster 1-4 maintained consistently short inter-residue distances of approximately 10 Å (8–13 Å) across all residue pairs, indicating sustained localization within the A*β* binding region.

To isolate meaningful interaction geometries from transiently dissociated states (such as the brief excursions observed late in the Cluster 1-4 trajectory), contact maps were rigorously filtered to include only bound-state frames where SKGQA resided within 5.0 Å of A*β*_17–25_ (Figure 7). To account for the intrinsically flexible nature of short peptides, these contact maps represent the thermodynamic spatial distribution of inter-residue contacts within the dynamic ensemble, rather than a single static complex geometry. Within these filtered bound ensembles, Cluster 1-4 exhibited remarkably high structural consistency across replicates (C*α* RMSD = 10.10 *±* 1.53 Å), confirming that its average contact distances reflect a well-defined, persistent binding mode. In contrast, Cluster 1-1 yielded a substantially higher C*α* RMSD (75.10 *±* 2.50 Å), indicating non-specific, transient spatial distributions where the peptide drifted away from the primary binding pocket. Remarkably, while Cluster 1-1 spent the vast majority of the simulation in a unbound/diffusive state (bound-state occupancy of only 4.1%), Cluster 1-4 maintained a remarkably high bound-state occupancy of 80.5%. This high population confirms that the late-stage RMSD fluctuations represent minor, reversible equilibrium events rather than permanent detachment, justifying the structural analysis focused on this robust bound ensemble. Within these bound ensembles, both clusters engaged the central A*β* segment with distinct spatial preferences. Specifically, Cluster 1-1 exhibited a binding distribution localized predominantly around the acidic region (Glu22–Asp23), where Gln4 made close contact with Glu22 (3.8 Å). Conversely, Cluster 1-4 shifted its binding interactions toward the central hydrophobic core (Phe20–Ala21), with Gln4 approaching Phe20 at 3.3 Å.

**Figure 7:**
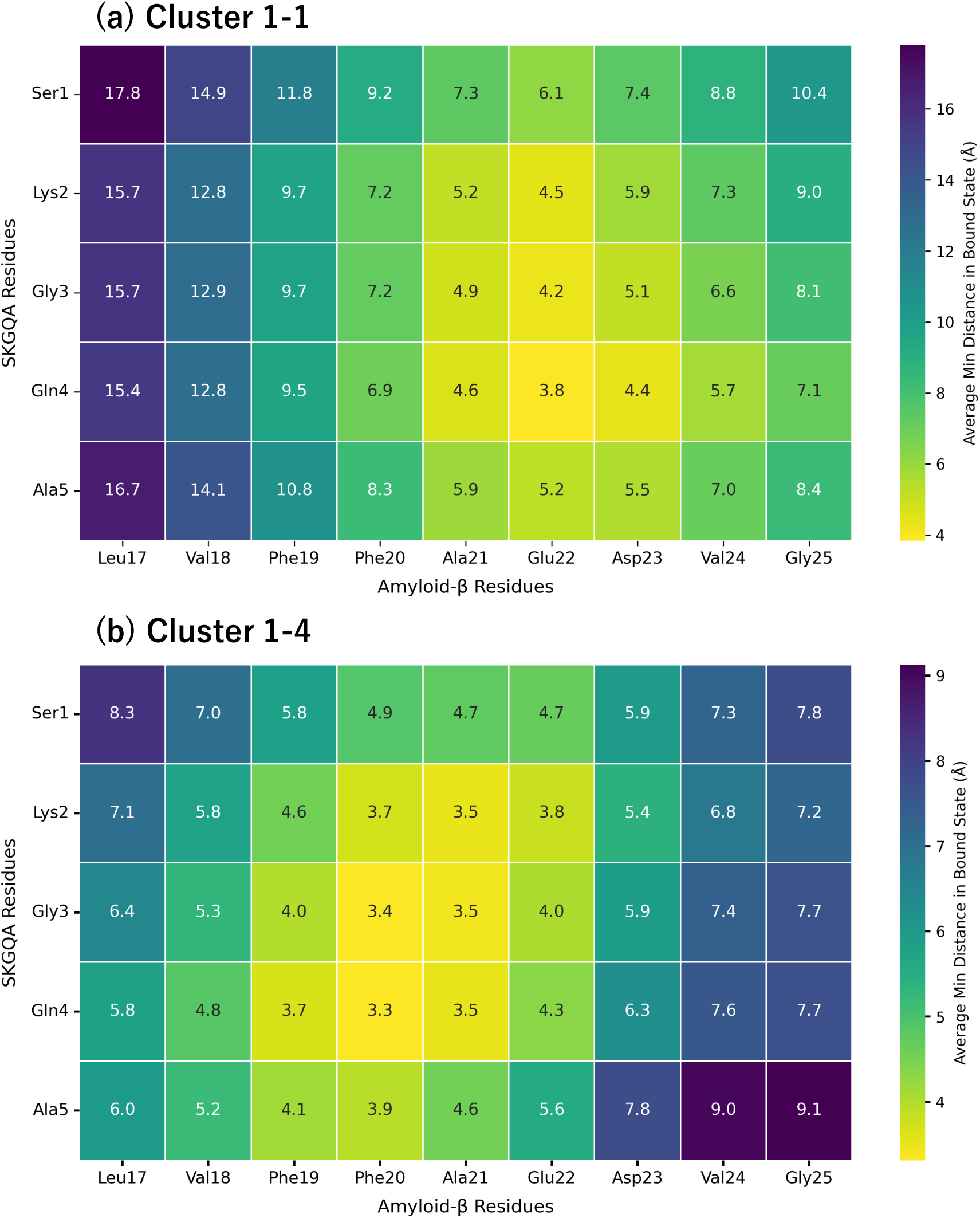
Residue-level contact maps of Cluster 1-1 and Cluster 1-4 in the bound state. Filtered contact maps show the average minimum interatomic distances (Å) between SKGQA residues (vertical axis: Ser1 to Ala5) and A*β* residues (horizontal axis: Leu17 to Gly25) for Cluster 1-1 (upper panel) and Cluster 1-4 (lower panel). Numeric values inside each cell indicate the exact average minimum distance (Å), with color scaling corresponding to spatial proximity as indicated by the color bar on the right.

To delineate the specific conformation when catalytic attack is feasible, a 4.0-Å filtered contact map (Reactive State) was generated, defined by the distance between the nucleophilic Ser1(O*γ*) of SKGQA and the backbone carbonyl carbons (*C* = O) of A*β* (Figure 8). Strikingly, while only 0.4% of frames in Cluster 1-1 met this reactive criterion compared to 31.4% in Cluster 1-4, both clusters formed highly convergent, productively oriented interaction interfaces upon entering the reactive state, while retaining their characteristic spatial biases.

**Figure 8:**
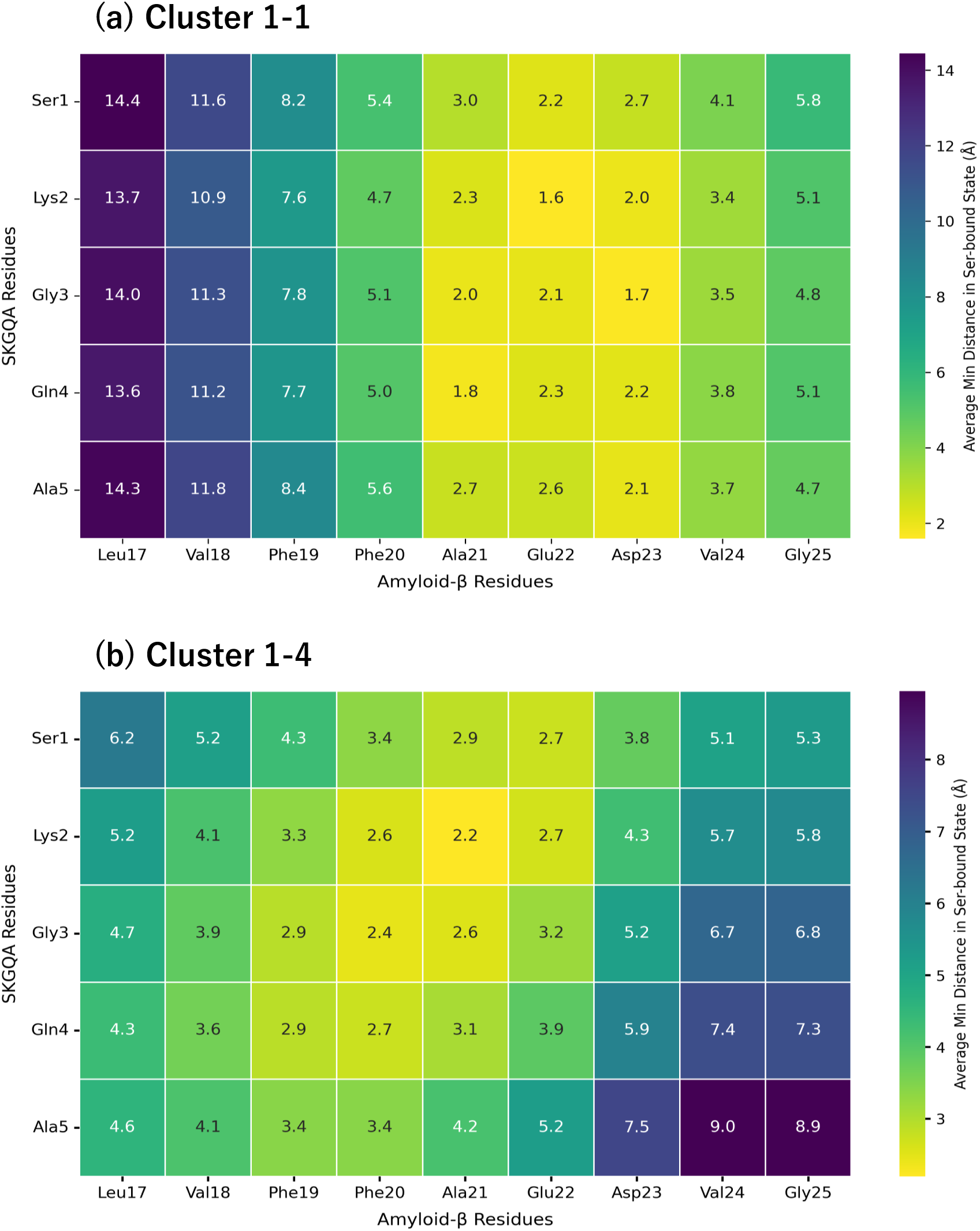
Residue-level contact maps of Cluster 1-1 and Cluster 1-4 in the reactive state. Filtered contact maps display the average minimum interatomic distances (Å) between SKGQA residues (vertical axis: Ser1 to Ala5) and A*β* residues (horizontal axis: Leu17 to Gly25) for Cluster 1-1 (upper panel) and Cluster 1-4 (lower panel), specifically calculated for frames in the reactive state (*≤* 4.0 Å distance between Ser1(O*γ*) and A*β* backbone carbonyl carbons). Numeric values inside each cell indicate the exact average minimum distance (Å), with color scaling corresponding to spatial proximity as indicated by the color bar on the right.

In Cluster 1-1 (Figure 8, upper panel), the reactive state was anchored around the acidic segment (Ala21–Asp23). Crucially, the catalytic Ser1 residue achieved minimal interatomic distances of 2.2 Å and 2.7 Å to Glu22 and Asp23, respectively, with Lys2 and Gly3 also making close approaches (*≤* 2.2 Å). In contrast, for Cluster 1-4 (Figure 8, lower panel), the reactive geometry was positioned further toward the central core (Phe20–Ala21). Here, Ser1 established close catalytic proximity to Ala21 (2.9 Å) and Glu22 (2.7 Å), supported by tight interactions from Lys2 (2.2 Å) and Gly3 (2.4 Å). These results demonstrate that although Cluster 1-1 binds transiently and Cluster 1-4 binds persistently, both clusters position the nucleophilic Ser1 residue within productive attack distance of the A*β* backbone, albeit targeting slightly shifted hotspots along the central segment (Ala21–Asp23 vs. Phe20–Ala21).

### 3.7 Structural Visualization and Atomic-Level Interaction of the Plausible Active Binding Mode

To elucidate the functional spatial orientation and atomic-level interactions of the bound state, a representative three-dimensional binding geometry was extracted by structural clustering (RMSD-based) of the bound-state trajectories originating from Cluster 1-4, which exhibited a dominant bound population (Figure 9).

**Figure 9:**
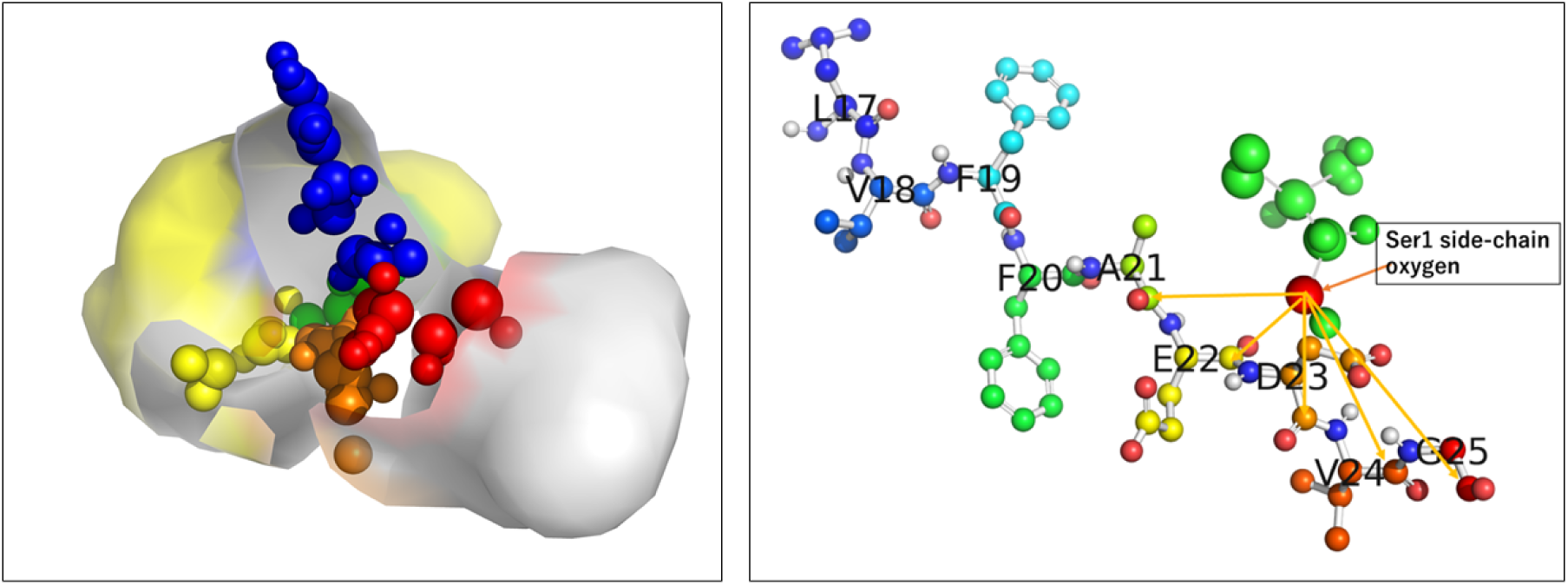
Representative three-dimensional binding geometry and atomic-level catalytic interactions of the SKGQA peptide in the bound state. **Left panel:** Global spatial distribution of the representative bound state of the SKGQA peptide on the A*β*_17–25_ surface, extracted by conformational clustering of bound trajectories from Cluster 1-4. SKGQA residues are represented as colored spheres: Ser1 (red), Lys2 (blue), Gly3 (green), Gln4 (orange), and Ala5 (yellow). The molecular surface of A*β*_17–25_ is color-coded by hydrophobicity, highlighting the central hydrophobic LVFFA core in yellow and the solvent-exposed hydrophilic/charged regions in white/gray. Ala5 acts as a hydrophobic anchor embedded within the LVFFA core, while Ser1 is positioned in the solvent-exposed region. **Right panel:** Atomic-level visualization (ball-and-stick model) of the reactive interface. The nucleophilic Ser1 side-chain oxygen atom (red sphere, indicated by label) is strategically oriented toward the backbone carbonyl carbons (*C* = O) of target A*β* residues within the cleavage zone (Ala21 through Gly25). Yellow arrows designate the directional paths for nucleophilic attack toward individual target carbonyl groups. The corresponding exact interatomic distances for each target site (from Leu17 to Gly25) are systematically summarized in Table 3.

**Table 3:** Interatomic distances between SKGQA Ser1(O*γ*) and backbone carbonyl carbons of A*β*_17*−*25_ in the representative reactive state.

| **Aβ Residue** | **Target Carbonyl Carbon** | **Distance (Å)** | **Attackability Evaluation** |
| --- | --- | --- | --- |
| Leu17 | Leu17 (C=O) | 9.38 | Distant |
| Val18 | Val18 (C=O) | 8.10 | Distant |
| Phe19 | Phe19 (C=O) | 7.64 | Out of range |
| Phe20 | Phe20 (C=O) | 6.19 | Intermediate |
| Ala21 | Ala21 (C=O) | 4.55 | Adjacent |
| Glu22 | Glu22 (C=O) | 2.78 | Highly plausible |
| Asp23 | Asp23 (C=O) | 2.35 | Highly plausible |
| Val24 | Val24 (C=O) | 3.08 | Adjacent |
| Gly25 | Gly25 (C=O) | 2.60 | Highly plausible |

The global spatial distribution (Figure 9, left panel) reveals a clear division of functional roles among the SKGQA residues upon binding to the A*β*_17–25_ surface: The C-terminal Ala5 residue (yellow spheres) embeds deeply into the central hydrophobic core (LVFFA, highlighted in yellow surface), functioning as a ”hydrophobic anchor” that firmly fixes the peptide positioning. Conversely, the catalytic Ser1 residue (red spheres) remains positioned toward the solvent-exposed hydrophilic outer region (white/gray surface), allowing its hydroxyl group (O*γ*) to freely engage the target backbone carbonyl carbons of A*β*. The intermediate residues—Lys2 (blue), Gly3 (green), and Gln4 (orange)—form a flexible bridging network that maintains solvent interactions and conformational adaptability. Remarkably, structural clustering of the rare reactive frames from Cluster 1-1 yielded an identical binding geometry to that of Cluster 1-4, confirming that this anchored orientation represents a thermodynamically favored, universal reactive state for A*β* cleavage.

To further detail the precise nucleophilic attack geometry, atomic-level interactions were visualized in ball-and-stick representation (Figure 9, right panel). The catalytic Ser1 side-chain oxygen (red sphere, labeled ”Ser1 side-chain oxygen”) directly faces the backbone carbonyl carbons (*C* = O) of the central A*β* segment (Ala21 to Gly25). Directional vectors (yellow arrows) highlight the catalytic accessibility from this nucleophilic oxygen toward multiple potential target sites.

To assess the feasibility of cleavage across multiple sites, interatomic distances between the Ser1(O*γ*) hydroxyl group and individual A*β*_17_*_−_*_25_ backbone carbonyl carbons (*C* = O) associated with these directional vectors were measured in this representative reactive structure (Table 3). The catalytic Ser1 side-chain oxygen is situated within immediate attack distance of multiple cleavage sites, particularly Glu22 (2.78 Å), Asp23 (2.35 Å), and Gly25 (2.60 Å), which are categorized as ”Highly Plausible” targets. Ala21 (4.55 Å) and Val24 (3.08 Å) reside in an ”Adjacent” zone, whereas N-terminal residues (Leu17–Phe19) are positioned farther away (*>* 7.5 Å). This spatial arrangement provides a structural rationale for the multi-site proteolytic cleavage observed experimentally at the A*β* central domain.

## 4 Discussion

### From Rigid Enzymes to Flexible Catalytides: A New Paradigm in Aβ Degradation

The development of small synthetic peptides, termed ”catalytides,” capable of degrading amyloid-beta (A*β*) represents a promising frontier in therapeutic design. We have previously demonstrated that the pentapeptide SKGQA, despite its minimal size, exhibits serine protease-like activity and cleaves A*β* at multiple sites[7]. However, a fundamental distinction exists between classical serine proteases and short catalytides. Natural serine proteases are large, rigid proteins evolved to maintain a pre-organized catalytic triad, enabling rapid and site-specific cleavage[8, 22]. In contrast, SKGQA operates over a much longer timescale and cleaves A*β* across multiple sites. In this study, we aimed to elucidate the dynamic molecular mechanism underlying this non-specific, multi-site cleavage using molecular dynamics simulations.

### Dynamic Conformational Diversity Drives Multi-site Cleavage

At the outset of this study, we hypothesized that SKGQA might adopt a stable, pre-organized active site mimicking natural serine proteases. However, our simulations demonstrate that short 5-mer catalytides operate through a fundamentally different paradigm: dynamic conformational diversity. Rather than locking into a single rigid structure, SKGQA retains high conformational freedom in solution and explores a heterogeneous binding ensemble upon encountering A*β*.

Crucially, this structural plasticity enables SKGQA to access distinct, functional interaction modes along the A*β* surface. Extended MD simulations revealed that SKGQA samples both a persistent binding mode (Cluster 1-4, occupying 80.5% of bound frames) and transient modes (such as Cluster 1-1). In Cluster 1-4, the C-terminal Ala5 acts as a hydrophobic anchor embedded within the central hydrophobic cluster (CHC, residues 17–21), while the catalytic Ser1 remains solvent-exposed near the hydrophilic/acidic region. This persistent geometry places Ser1(O*γ*) within immediate proximity of target backbone carbonyl carbons, particularly Glu22 (2.78 Å) and Asp23 (2.35 Å). These spatial proximity values align well with the experimental cleavage sites identified in Figure 2. Although Ser1 also approaches Asp23 closely in the simulations, cleavage at Asp23 was not experimentally observed. This distinction highlights that while atomic proximity is a prerequisite for nucleophilic attack, actual proteolysis is ultimately governed by intrinsic chemical reactivity and cleavage kinetics. Overall, the ability of SKGQA to explore multiple reactive geometries across both hydrophobic and acidic microenvironments provides a clear structural rationalization for its experimentally observed multi-site cleavage.

### Structural Flexibility and Dynamic Ensemble Nature of the Minimal 5-mer Peptide

It is important to emphasize that short peptides such as SKGQA naturally lack a rigid, pre-organized three-dimensional tertiary structure in aqueous solution. Unlike bulky globular enzymes that rely on a rigid active site scaffold, the minimal 5-mer peptide exhibits intrinsically high conformational flexibility. Consequently, its functional mechanism relies on a dynamic equilibrium—encompassing conformational selection and substrate-induced fit—rather than binding to a single, static energy-minimum state [23]. Throughout the molecular dynamics (MD) trajectories, the persistent fluctuation observed in the RMSD and inter-atomic distances does not represent a lack of convergence or system instability. Instead, it reflects the true physical behavior of the short peptide exploring its conformational energy landscape. Transient activation occurs when thermal motion periodically brings the catalytic residues into a favorable geometry close to the A*β* backbone. Therefore, evaluating this catalytic system requires capturing the ensemble distribution of these transient, reactive states rather than searching for a rigid, permanent binding pose, aligned with established principles of catalytic peptide assemblies [24, 25].

### Interference with Aβ Self-Assembly and Amyloid Cascade Strategy

Beyond targeted proteolysis, the interaction of SKGQA with the central hydrophobic core (CHC; Aβ_17–21_) region has critical implications for Aβ aggregation. The CHC is well recognized as an essential recognition motif driving Aβ nucleation, oligomerization, and fibril assembly [26, 27]. By embedding its hydrophobic Ala5 residue into the CHC interface, SKGQA may act as a physical inhibitor that sterically blocks self-assembly, consistent with CHC-targeted peptide inhibition strategies [27, 28].

According to the classical amyloid cascade hypothesis, Aβ transitions dynamically through a monomer–oligomer–fibril equilibrium (Monomer ⇌ Oligomer ⇌ Fibril) [29, 30]. Because soluble monomers and toxic oligomers exist in a dynamic equilibrium, degradation of monomeric Aβ serves as an effective upstream strategy to shift the equilibrium away from oligomerization, thereby reducing toxic aggregate formation. Furthermore, the amphiphilic balance of SKGQA (combining hydrophobic anchoring via Ala5 and electrostatic guidance via Lys2/Glu22) prevents irreversible entrapment, allowing the peptide to scan aggregate surfaces and potentially disaggregate pre-formed fibrils.

### Study Limitations and Future Perspectives

While this study offers critical mechanistic insights into catalytide dynamics, several methodological limitations warrant consideration. Regarding initial conformation and target region selection, a single representative SKGQA conformation—selected based on a short N–C terminal distance—was used for initial docking via HADDOCK, with active residues restricted specifically to the CHC region (LVFFA, A*β*_17–21_). Because experimental data demonstrate that SKGQA cleaves A*β* at multiple sites beyond the CHC, simulating alternative initial SKGQA conformations and targeting other regions across the A*β* sequence will be essential to fully elucidate the complete landscape of multi-site cleavage. Additionally, our current MD simulations focused on monomeric monomeric A*β*_17–42_; further studies utilizing oligomeric or fibrillar A*β* assemblies are needed to directly evaluate the structural disaggregation and cleavage mechanisms in higher-order aggregate states.

### Clinical Potential Beyond Antibody Therapies

Compared to monoclonal antibody therapies (e.g., lecanemab)[31], which target specific aggregate surfaces and suffer from limited brain penetration, small catalytic peptides offer significant advantages in tissue permeability, delivery, and cost-effectiveness. Recent *in vivo* studies on related catalytic peptides (catalytides) provide an encouraging proof-of-concept for their therapeutic viability; for instance, the short catalytide GSGFK was shown to rescue short-term memory deficits in A*β*_25_*_−_*_35_-induced Alzheimer’s mouse models by suppressing and reversing aggregation[32, 33]. Although A*β*_25_*_−_*_35_ represents a specific truncated substrate fragment distinct from our CHC-targeted (A*β*_17_*_−_*_21_) model, these findings demonstrate that minimal catalytides can retain catalytic stability and functional efficacy *in vivo*. Additionally, non-invasive intranasal (i.n.) delivery of short catalytides (such as ANA-TA9) achieves efficient direct brain distribution via the olfactory epithelium, providing an ideal, non-invasive platform for long-term preventive therapy[34]. Optimizing SKGQA sequence variants and delivery routes holds immense therapeutic potential for Alzheimer’s disease and broader amyloid disorders.

## 5 Conclusions

Our combined computational and experimental findings demonstrate that the minimal 5-mer catalytide SKGQA executes multi-site cleavage of A*β* through a flexible, substrate-assisted catalytic mechanism rather than a rigid catalytic triad. Hydrophobic anchoring of the C-terminal Ala5 to the A*β* CHC region (residues 17–21) stabilizes the enzyme–substrate complex, positioning the hydrophilic N-terminal Ser1 in spatial proximity to multiple adjacent backbone carbonyl carbons, including Glu22 and Asp23. Within this anchored conformational ensemble, thermal fluctuations enable Ser1 to stochastically attack these neighboring target sites, providing a clear structural rationale for the peptide’s multi-site proteolytic activity. Overall, these insights confirm that small, highly flexible peptides can achieve efficient catalytic function, offering a promising foundation for the rational design of cost-effective, site-specific peptide therapeutics for Alzheimer’s disease.

## 6 Patents

AMYLOID-*β* AGGREGATION INHIBITOR, PHARMACEUTICAL COMPOSITION FOR AMYLOID-*β* AGGREGATION DISEASES, AND USE APPLICATION OF SAME, WO/2022/239765. However, this patent was not derived from the current simulation study.

## Statements and Declarations

### Author Contributions

Conceptualization, F.I. and T.A.; research planning, analyses or interpretation of data, F.I.; writing—original draft preparation, F.I.; writing—review and editing, T.A., M.K. and R.N. All authors have read and agreed to the published version of the manuscript.

### Institutional Review and Informed Consent

Consent for institutional review was not required, and informed consent was not applicable to this study.

### Data Availability

The data used in this study are available from the corresponding authors upon reasonable request.

## Acknowledgments

During the preparation of this work, the author used Gemini to refine the English expression and improve the logical clarity of the manuscript content. The author takes full responsibility for the scientific integrity and content of the publication.

## Abbreviations

The following abbreviations are used in this manuscript:

AD: Alzheimer’s disease A*β* amyloid-*β*
MD: molecular dynamics
RMSD: Root Mean Square Deviation
CHC: Central Hydrophobic Cluster

## References

[1] Mary P. Lambert, Ann K. Barlow, Brett A. Chromy, Carrie Berry, William A. Frost, Geoffrey A. Higgins, Grant A. Krafft, and William L. Klein. Diffusible, non-fibrillar ligands derived from A*β*_1*−*42_ are potent central nervous system neurotoxins. Proceedings of the National Academy of Sciences of the United States of America, 95(11):6448–6453, 1998.

[2] Urmi Sengupta, Ashley N. Nilson, and Rakez Kayed. The role of amyloid-*β* oligomers in toxicity, propagation, and immunotherapy. EBioMedicine, 6:42–49, 2016.

[3] Eric McDade, Jeffrey L. Cummings, Shobha Dhadda, Chad J. Swanson, Larisa Reyderman, Michio Kanekiyo, Akihiko Koyama, Michael Irizarry, Lynn D. Kramer, and Randall J. Bateman. Lecanemab in patients with early Alzheimer’s disease: detailed results on biomarker, cognitive, and clinical effects from the randomized and open-label extension of the phase 2 proof-of-concept study. Alzheimer’s Research & Therapy, 14(1):191, 2022.

[4] Sarah J. Doran and Russell P. Sawyer. Risk factors in developing amyloid related imaging abnormalities (ARIA) and clinical implications. Frontiers in Neuroscience, 18:1337029, 2024.

[5] Rina Nakamura, Motomi Konishi, Youichirou Higashi, Motoaki Saito, and Toshifumi Akizawa. Comparison of the catalytic activities of 5-mer synthetic peptides derived from box A region of Tob/BTG family proteins against the amyloid-beta fragment peptides. Integrative Molecular Medicine, 6:1–6, 2019.

[6] Rina Nakamura, Motomi Konishi, Masanari Taniguchi, Yusuke Hatakawa, and Toshifumi Akizawa. The discovery of shorter synthetic proteolytic peptides derived from Tob1 protein. Peptides, 116:71–77, 2019.

[7] Yusuke Hatakawa, Rina Nakamura, Toshifumi Akizawa, Motomi Konishi, Akira Matsuda, Tomoyuki Oe, Motoaki Saito, and Fumiaki Ito. SKGQA, a peptide derived from the ANA/BTG3 protein, cleaves amyloid-*β* with proteolytic activity. Biomolecules, 14(5):568, 2024.

[8] Lizbeth Hedstrom. Serine protease mechanism and specificity. Chemical Reviews, 102(12):4501–4524, 2002.

[9] Alexis Lamiable, Pierre Thévenet, Julien Rey, Marek Vavruška, Philippe Derreumaux, and Pierre Tufféry. PEP-FOLD3: faster de novo structure prediction for linear peptides in solution and in complex. Nucleic Acids Research, 44(W1):W449– W454, 2016.

[10] Pierre Thévenet, Yimin Shen, Julien Maupetit, Fŕedéric Guyon, Philippe Derreumaux, and Pierre Tufféry. PEP-FOLD: An updated de novo structure prediction server for both linear and disulfide bonded cyclic peptides. Nucleic Acids Research, 40(W1):W288–W293, 2012.

[11] Julien Maupetit, Yimin Shen, Pierre Tufféry, and Philippe Derreumaux. Improved PEP-FOLD approach for peptide and miniprotein structure prediction. Journal of Chemical Theory and Computation, 10(10):4745–4758, 2014.

[12] Benjamin Webb and Andrej Sali. Comparative protein structure modeling using MODELLER. Current Protocols in Bioinformatics, 54(1):5.6.1–5.6.37, 2016.

[13] Schrödinger, LLC and Warren L. DeLano. The PyMOL molecular graphics system, version 2.0, 2020.

[14] Thorsten Lührs, Christiane Ritter, Marc Adrian, Dominique Riek-Loher, Bernd Bohrmann, Heinz Döbeli, David Schubert, and Roland Riek. 3D structure of Alzheimer’s amyloid-*β*(1 *−* 42) fibrils. Proceedings of the National Academy of Sciences of the United States of America, 102(48):17342–17347, 2005.

[15] Rodrigo V. Honorato, Panagiotis I. Koukos, Brian Jiménez-Garćıa, Andrei Tsare-gorodtsev, Marco Verlato, Andrea Giachetti, Antonio Rosato, and Alexandre M. J. J. Bonvin. Structural biology in the clouds: The WeNMR-EOSC ecosystem. Frontiers in Molecular Biosciences, 8:712926, 2021.

[16] Rodrigo V. Honorato, Mikael E. Trellet, Brian Jiménez-Garćıa, Jörg J. Schaarschmidt, Marco Giulini, Victor Reys, Panagiotis I. Koukos, Jõao P. G. L. M. Rodrigues, Ezgi Karaca, Gydo C. P. van Zundert, Jorge Roel-Touris, Charlotte W. van Noort, Zuzana Jandová, Adrien S. J. Melquiond, and Alexandre M. J. J. Bonvin. The HADDOCK2.4 web server for integrative modeling of biomolecular complexes. Nature Protocols, 19(11):3219–3241, 2024.

[17] Peter Eastman, Raimondas Galvelis, Raúl P. Peĺaez, Charlles R. A. Abreu, Stephen E. Farr, Emilio Gallicchio, Anton Gorenko, Michael M. Henry, Frank Hu, Jing Huang, Andreas Krämer, Julien Michel, Joshua A. Mitchell, Vijay S. Pande, João P. G. L. M. Rodrigues, Jaime Rodriguez-Guerra, Andrew C. Simmonett, Sukrit Singh, Jason Swails, Philip Turner, Yuanqing Wang, Ivy Zhang, John D. Chodera, Gianni De Fabritiis, and Thomas E. Markland. OpenMM 8: Molecular dynamics simulation with machine learning potentials. The Journal of Physical Chemistry B, 128(1):109–116, 2024.

[18] James A. Maier, Channing Martinez, Koushik Kasavajhala, Lauren Wickstrom, Amy K. E. Hauser, and Carlos Simmerling. ff14SB: Improving the accuracy of protein side chain and backbone parameters from ff99SB. Journal of Chemical Theory and Computation, 11(8):3696–3713, 2015.

[19] William L. Jorgensen, Jayaraman Chandrasekhar, Jeffry D. Madura, Roger W. Impey, and Michael L. Klein. Comparison of simple potential functions for simulating liquid water. The Journal of Chemical Physics, 79(2):926–935, 1983.

[20] Ulrich Essmann, Lalith Perera, Max L. Berkowitz, Tom Darden, Hsing Lee, and Lee G. Pedersen. A smooth particle mesh Ewald method. The Journal of Chemical Physics, 103(19):8577–8593, 1995.

[21] Robert T. McGibbon, Carlos X. Hernandez, Matthew P. Harrigan, Steven Kearnes, Mohammad M. Sultan, Sian Kobutzke, Christoph Wehmeyer, Jan Bauer, and Vijay S. Pande. MDTraj: A modern open library for the analysis of molecular dynamics trajectories. Biophysical Journal, 109(8):1528–1532, 2015.

[22] Özlem Doğan Ekici, Mark Paetzel, and Ross E. Dalbey. Unconventional serine proteases: Variations on the catalytic Ser/His/Asp triad configuration. Protein Science, 17(12):2023–2037, 2008.

[23] David D. Boehr, Ruth Nussinov, and Peter E. Wright. The role of dynamic conformational ensembles in biomolecular recognition. Nature Chemical Biology, 5(11):789– 796, 2009.

[24] Scott J. Miller. In search of peptide-based catalysts for asymmetric organic synthesis. Accounts of Chemical Research, 37(8):601–610, 2004.

[25] Jefferson D. Revell and Helma Wennemers. Peptidic catalysts developed by combinatorial screening methods. Current Opinion in Chemical Biology, 11(3):269–278, 2007.

[26] William P. Esler, Evelyn R. Stimson, Jennifer M. Jennings, Harry V. Vinters, Joseph R. Ghilardi, Patrick W. Mantyh, and John E. Maggio. Point mutations and chemical modifications in A*β*(10 *−* 35) identify the hydrophobic core as essential for amyloid assembly and deposition. Biochemistry, 35(44):13914–13921, 1996.

[27] Lars O. Tjernberg, Jan Näslund, Fredrik Lindqvist, Jan Johansson, A. Rickard Karlström, Johan Thyberg, Lars Terenius, and Charles Nordstedt. Arrest of *β*-amyloid fibril formation by a pentapeptide ligand. Journal of Biological Chemistry, 271(15):8545–8548, 1996.

[28] John R. Horsley, Blagojce Jovcevski, Kate L. Wegener, Jingxian Zhang, Xiaocong Qiu, Steven W. Polyak, John A. Carver, and Andrew D. Abell. Peptide inhibitors targeting the amyloid-*β* self-recognition sequence KLVFF prevent aggregation and toxicity. Biochemical Journal, 477(11):2039–2039, 2020.

[29] David B. Teplow, Nicholas D. Lazo, Gal Bitan, Summer L. Bernstein, Thomas Wyttenbach, Michael T. Bowers, Sabina S. Vollers, Margaret M. Condron, Brian A. Reed, and M. Melody Lin. Amyloid *β*-protein assembly and Alzheimer disease. Journal of Biological Chemistry, 284(8):4749–4753, 2009.

[30] Subrata Nag, Bipasha Sarkar, Ananya Bandyopadhyay, Bikash R. Sahoo, V. K. A. Sreenivasan, Mindy Kombrabail, C. N. R. Rao, and Sudipta Maiti. Nature of the amyloid-*β* monomer and the monomer-oligomer equilibrium. Journal of Biological Chemistry, 286(16):13827–13833, 2011.

[31] Chad J. Swanson, Yong Zhang, Shobha Dhadda, Jinping Wang, June Kaplow, Robert Y. K. Lai, Lars Lannfelt, Heather Bradley, Martin Rabe, Akihiko Koyama, Larisa Reyderman, Donald A. Berry, Scott Berry, Robert Gordon, Lynn D. Kramer, and Jeffrey L. Cummings. A randomized, double-blind, phase 2b proof-of-concept clinical trial in early Alzheimer’s disease with lecanemab, an anti-A*β* protofibril antibody. Alzheimer’s Research & Therapy, 13(1):80, 2021.

[32] Rina Nakamura, Motomi Konishi, Youichirou Higashi, Motoaki Saito, and Toshifumi Akizawa. Five-mer peptides prevent short-term spatial memory deficits in A*β*_25*−*35_-induced Alzheimer’s model mouse by suppressing A*β*_25*−*35_ aggregation and resolving its aggregate form. Alzheimer’s Research & Therapy, 15(1):83, 2023.

[33] Rina Nakamura, Akira Matsuda, Youichirou Higashi, Yoshihiro Hayashi, Motomi Konishi, Motoaki Saito, and Toshifumi Akizawa. An 11-mer synthetic peptide suppressing aggregation of A*β*_25*−*35_ and resolving its aggregated form improves test performance in an A*β*_25*−*35_-induced Alzheimer’s mouse model. Biomolecules, 14(11):1364, 2024.

[34] Yusuke Hatakawa, Akiko Tanaka, Tomoyuki Furubayashi, Rina Nakamura, Motomi Konishi, Toshifumi Akizawa, and Toshiyasu Sakane. Direct delivery of ANA-TA9, a peptide capable of A*β* hydrolysis, to the brain by intranasal administration. Pharmaceutics, 13(11):1832, 2021.

