## Supplementary Figures and Tables for "Multi-site Cleavage of Amyloid-β by a Minimal 5-mer Catalytic Peptide: Mimicking Serine Protease Activity via Hydrophobic Anchoring"

### Supplementary Information

### Supplementary Figures

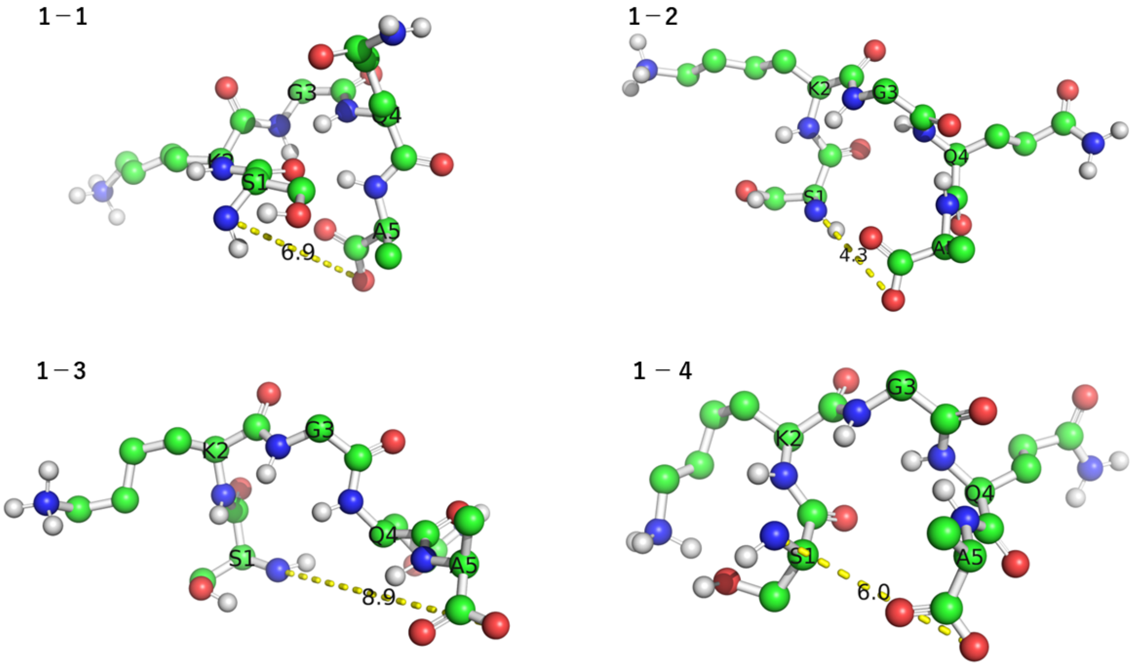


Figure S1: **Bound conformations and intramolecular Ser1(N)–Ala5(O/OXT) distances of SKGQA models in Cluster 1.** Three-dimensional structures of bound SKGQA peptides extracted from the four docking models of Cluster 1 (Clusters 1-1, 1-2, 1-3, and 1-4). Amino acid residues (Ser1 to Ala5, labeled as S1–A5) are shown in ball-and-stick representations. Yellow dashed lines and corresponding numbers indicate the intramolecular distance (Å) between the N-terminal nitrogen atom of Ser1 and the C-terminal carboxylate oxygen of Ala5, highlighting the substantial conformational variability of bound SKGQA across different docking poses (ranging from 4.3 to 8.9 Å).Description and details of Figure S1.

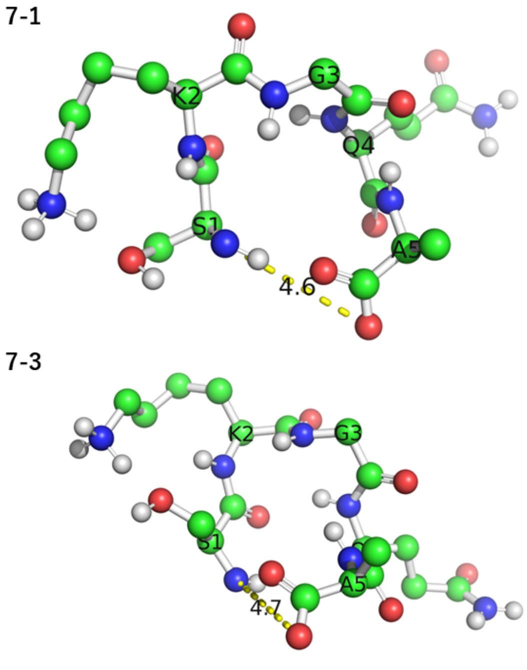


Figure S2: **Bound conformations and intramolecular Ser1(N)–Ala5(O/OXT) distances of SKGQA models in Cluster 7.** Three-dimensional structures of bound SKGQA peptides extracted from the four docking models of Cluster 7 (Clusters 7-1, 7-2, 7-3, and 7-4). Amino acid residues (Ser1 to Ala5, labeled as S1–A5) are shown in ball-and-stick representations. Yellow dashed lines and corresponding numbers indicate the intramolecular distance (Å) between the N-terminal nitrogen atom of Ser1 and the C-terminal carboxylate oxygen of Ala5, illustrating conformational flexibility across Cluster 7 models (ranging from 4.6 to 8.5 Å).Description and details of Figure S1.

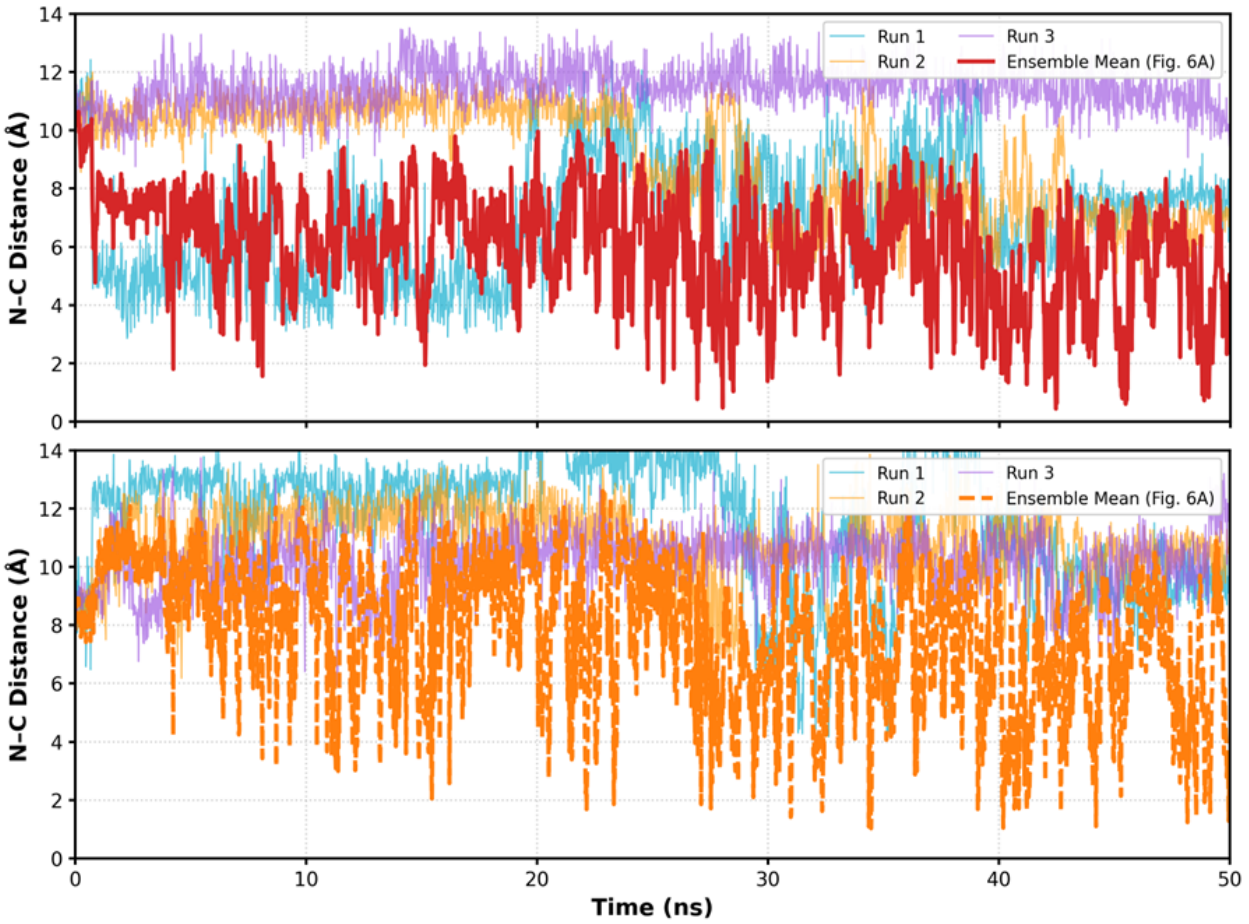


Figure S3: **Single-run trajectory profiles and ensemble averages of N–C distances for Cluster 1-1 over 50-ns MD simulations.** Time-dependent intramolecular N–C distance profiles (Å) within SKGQA for Cluster 1-1 over 0–50 ns, comparing three independent production trajectories (Run 1: cyan, Run 2: yellow, and Run 3: purple) alongside the overall ensemble mean profile across the replicate simulations. Upper panel: Distances between Ser1(N) and the terminal carboxylate oxygen Ala5(O1), with the ensemble mean shown as a solid red line (as presented in Fig. 6a). Lower panel: Distances between Ser1(N) and Ala5(O2), with the ensemble mean shown as a dashed orange line (as presented in Fig. 6a). Individual runs exhibit distinct temporal fluctuations, highlighting the diverse conformational space explored across independent replicates.

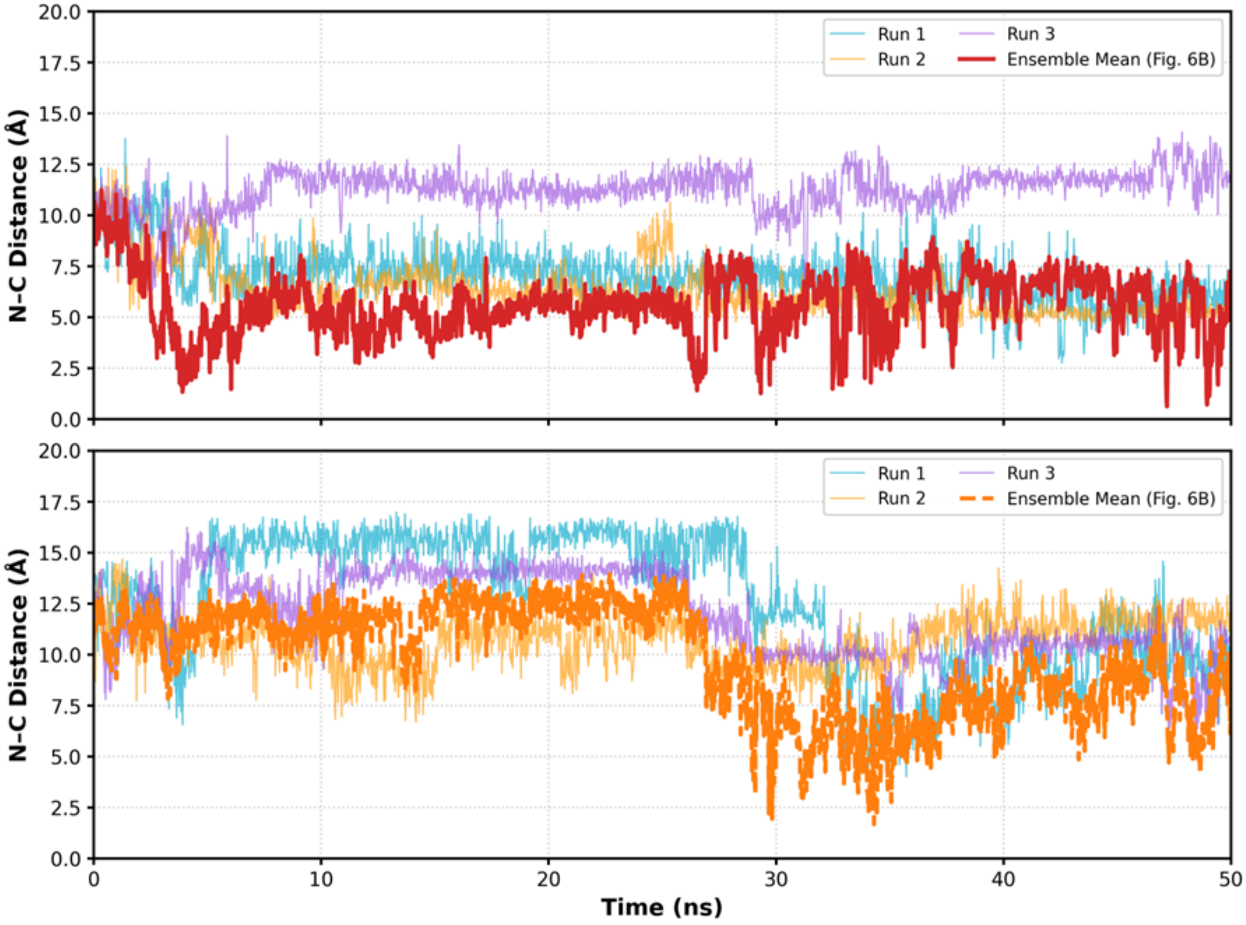


Figure S4: **Single-run trajectory profiles and ensemble averages of N–C distances for Cluster 1-4 over 50-ns MD simulations.** Time-dependent intramolecular N–C distance profiles (Å) within SKGQA for Cluster 1-4 over 0–50 ns, comparing three independent production trajectories (Run 1: cyan, Run 2: yellow, and Run 3: purple) alongside the overall ensemble mean profile across the replicate simulations. Upper panel: Distances between Ser1(N) and the terminal carboxylate oxygen Ala5(O1), with the ensemble mean shown as a solid red line (as presented in Fig. 6b). Lower panel: Distances between Ser1(N) and Ala5(O2), with the ensemble mean shown as a dashed orange line (as presented in Fig. 6b). Individual runs exhibit distinct temporal fluctuations, highlighting the diverse conformational sampling explored across independent replicates.

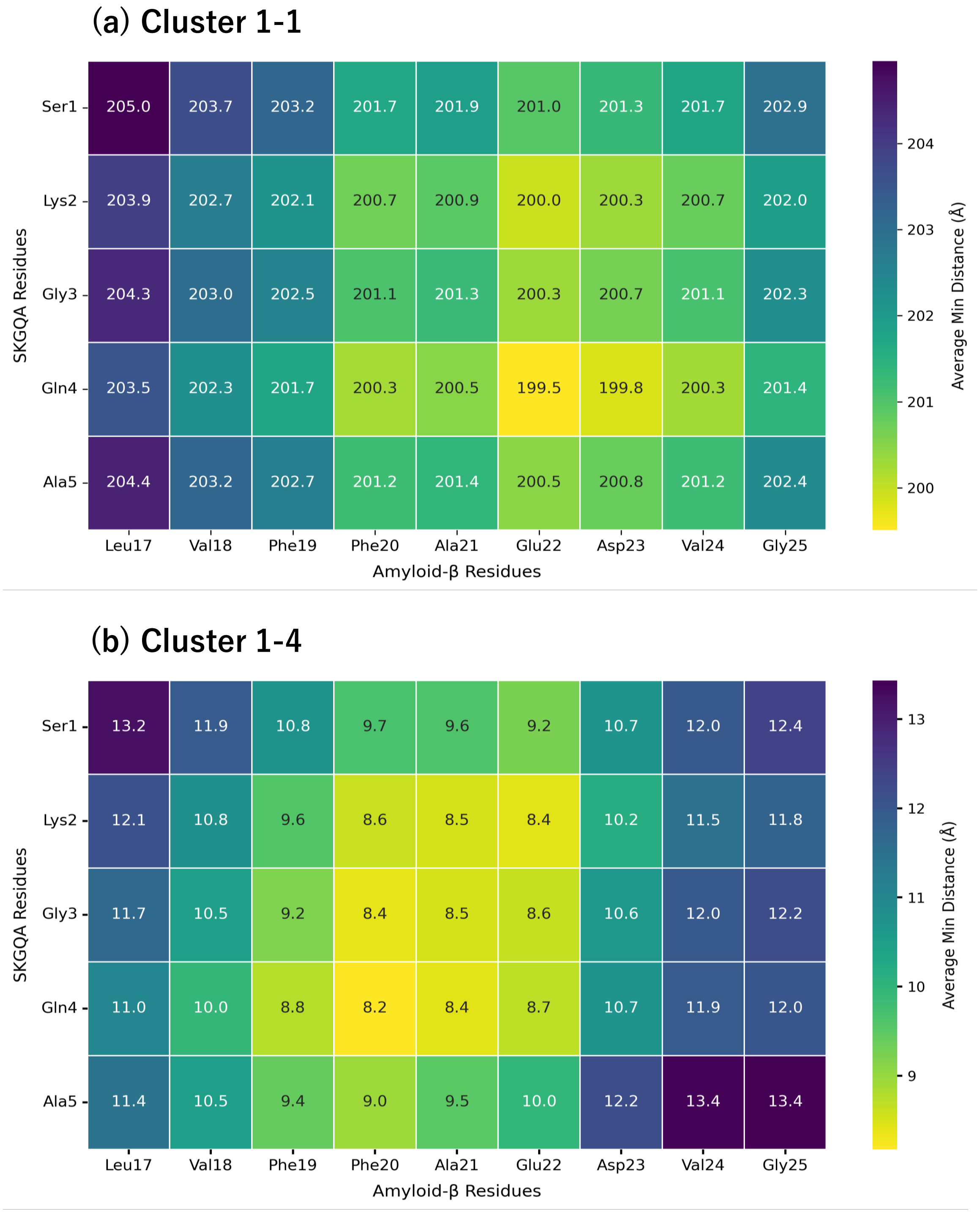


Figure S5: **Inter-residue average minimum distance contact maps for Cluster 1-1 and Cluster 1-4 over 50 ns MD simulations.** Residue-level average minimum distance maps (Å) evaluated across all frames of the 50-ns simulation trajectories between SKGQA residues (Ser1 to Ala5) and A$\beta_{17\text{–}25}$ residues (Leu17 to Gly25). **(a) Cluster 1-1:** Average minimum distances exceed 200 Å (ranging from $\sim$ 199.5 to 205.0 Å), indicating a fully dissociated state where SKGQA diffuses away from A$\beta_{17\text{–}25}$. **(b) Cluster 1-4:** Average minimum distances remain consistently short (ranging from 8.2 to 13.4 Å), demonstrating persistent spatial localization and stable contact with the central region of A$\beta_{17\text{–}25}$.
